# Phenylketonuria: modelling cerebral amino acid and neurotransmitter metabolism

**DOI:** 10.1101/2024.01.05.574352

**Authors:** Agnieszka B. Wegrzyn, Danique van Vliet, Karen van Eunen, M. Rebecca Heiner-Fokkema, Eddy A. van der Zee, Francjan J. van Spronsen, Barbara M. Bakker

**Author notes:** equal contributions (Agnieszka B. Wegrzyn, Danique van Vliet, Francjan J. van Spronsen, Barbara M. Bakker).

## Abstract

**Objective:** Phenylketonuria (PKU) is a metabolic disorder characterised by deficient hepatic phenylalanine hydroxylase activity, leading to elevated phenylalanine levels. Despite adherence to a phenylalanine-restricted diet, many adult PKU patients continue to experience executive function deficits, likely linked to high cerebral phenylalanine concentrations and deficiencies in monoaminergic neurotransmitters. Given the complexity of the interaction between diet and brain neurotransmitter metabolism, we employed computational metabolic modelling alongside experimental data from dietary intervention studies in PKU mice to identify key metabolic drivers underlying these deficits. Our goal was to provide a mechanistic, model-based foundation to support and optimise dietary therapies in PKU.

**Method:** We developed a computational model simulating large neutral amino acid (LNAA) transport across the blood-brain barrier and the subsequent metabolism of cerebral amino acids and monoaminergic neurotransmitters. The model was validated using direct measurements of brain amino acid concentrations in PKU mice subjected to various dietary regimens.

**Results:** The model predicts that cerebral amino acid levels are primarily influenced by their plasma concentrations and, to a lesser extent, by competition among LNAAs for transport mechanisms. Notably, it suggests that cerebral monoaminergic neurotransmitter levels are more significantly affected by elevated phenylalanine levels, likely through non-competitive inhibition of hydroxylase enzymes, than by the availability of precursor amino acids. Consequently, the model indicates that reducing phenylalanine levels, in conjunction with supplementing tyrosine and tryptophan, is more effective in restoring neurotransmitter levels than precursor supplementation alone.

**Conclusion:** This study presents the first comprehensive model integrating LNAA transport and cerebral neurotransmitter metabolism in PKU. The model enhances our understanding of the disease’s pathophysiology and highlights the importance of combined therapeutic strategies that target both phenylalanine reduction and precursor amino acid supplementation. Furthermore, it identifies knowledge gaps in LNAA transport mechanisms and offers a framework applicable to other neurological disorders involving diet-gene-neurotransmitter interactions.

## Introduction

Phenylketonuria (PKU; OMIM 261600) is the classic example of an inborn error of amino acid metabolism. It is caused by a deficiency of the hepatic phenylalanine hydroxylase (PAH), which converts phenylalanine into tyrosine [1]. As the distinguishment between ‘PKU’, ’classical PKU’ and other forms of PAH deficiency is rather unclear, we here refer to PAH deficiency causing cerebral pathophysiology referred to as ‘PKU’. If left untreated, high plasma phenylalanine rather than low tyrosine concentrations have been associated with PKU symptomatology. The latter is almost exclusively restricted to brain functioning, including severe intellectual disability, seizures, motor deficits, and psychiatric problems. Today, neonatal screening allows PKU diagnosis and initiation of treatment shortly after birth. The cornerstone of the treatment is to reduce phenylalanine concentrations in blood and brain by a severe phenylalanine-restricted diet. This diet consists of three parts: 1) a diet very low in natural protein; 2) a protein substitute, supplementing all amino acids except phenylalanine (and tyrosine-enriched) and other micronutrients that are normally present in high natural protein containing food, and 3) low-protein food (that especially supplies patients with energy) [2,3].

Additionally, some patients respond to tetrahydrobiopterin supplementation. Tetrahydrobiopterin is a natural co-substrate of hepatic PAH, and of (cerebral) tyrosine and tryptophan hydroxylases. In addition, to being their redox co-substrate, it may act as a pharmaceutical chaperone of PAH, supporting its conformational stability and preventing degradation [3,4]. Furthermore, in 2018 FDA has approved an injectable pegylated phenylalanine ammonia lyase, which can lower the phenylalanine levels in most patients. However, the leads to several, difficult to manage side effects [3]. While the phenylalanine-restricted diet and existing pharmacological treatments can prevent severe intellectual disability [5], the clinical outcome remains suboptimal and warrants additional/alternative pathophysiology-based treatment strategies [1].

In PKU pathophysiology, the blood-brain barrier (BBB) is considered to play a central role [6,7]. Phenylalanine, as well as all other large neutral amino acids (LNAA), such as tyrosine and tryptophan, are exchanged across the BBB by the large neutral amino acid transporter 1 (LAT1) [8]. This causes competition between the different LNAA for LAT1. Consequently, excessive plasma phenylalanine concentrations may not only lead to increased brain phenylalanine levels but also outcompete the transport of other LNAA across the BBB and thereby impair their brain availability [9–11]. While high brain phenylalanine levels are neurotoxic and affect brain metabolism [12–16], insufficient brain availability of non-phenylalanine LNAA has been related to impaired cerebral protein synthesis [11,17]. Moreover, tyrosine and tryptophan are the precursors for the cerebral monoaminergic neurotransmitters dopamine, norepinephrine, and serotonin, respectively [18]. High cerebral phenylalanine content is known to inhibit (cerebral) tyrosine and tryptophan hydroxylases, which are the enzymes performing the rate-limiting steps in the synthesis of dopamine and serotonin [19–22]. Thus, a combination of high phenylalanine and low tyrosine and tryptophan may lead to low concentrations of monoaminergic neurotransmitters. This has been suggested to play an important role in the mood and psychosocial problems of PKU patients [23–25]. However, while phenylalanine neurotoxicity, impaired cerebral protein synthesis and reduced cerebral monoaminergic neurotransmitter synthesis have all been associated with brain dysfunction in PKU patients [6]. Both cerebral protein synthesis and cerebral monoaminergic neurotransmitter synthesis show correlations with the plasma phenylalanine concentration that comprises the main treatment target and biomarker in today’s PKU management.

Based on this pathophysiological concept, supplementation of non-phenylalanine LNAA instead of restricting dietary phenylalanine intake has been suggested as a possible alternative non-pharmaceutical treatment strategy [6,26]. Such LNAA treatment has been shown in PKU mice to (1) reduce brain phenylalanine, (2) increase brain non-phenylalanine LNAA, and (3) increase brain monoaminergic neurotransmitter concentrations [27]. To ultimately establish the adequate dose of different LNAA, a better understanding of the pathophysiology of brain dysfunction in PKU, and especially of the relationships between plasma and brain amino acid concentrations, and between these amino acid and monoamine concentrations, is essential.

The competition of amino acids for the LAT1 transporter and the involvement of additional amino acid transporters complicate the understanding of the effect of different diets on the brain amino acid and neurotransmitter composition. Such complex systems can be studied holistically using computational models. Two distinct modelling approaches exist that allow mechanistic studies of metabolic networks: 1) genome-scale constraint-based models, and 2) kinetic, ordinary differential equation (ODE)-type models. The genome-scale models comprise the entirety of the known metabolic network, including reaction stoichiometry and mass and charge balance. However, due to their scale and in contrast to kinetic models, they lack kinetic information and regulation [28]. Kinetic models usually represent only smaller pathways, due to their increased mechanical complexity and difficulty of obtaining accurate kinetic parameters [29].

To address the problem of substrate competition for LAT1 amino acid transport in PKU, several computational models have been constructed that describe the kinetic behaviour of pathways involved in neurotransmitter metabolism or LNAA transport across the BBB [30–35]. However, none of these models integrates the dopaminergic and serotonergic pathways in the brain with the amino acid transport across the BBB. Constraint-based stoichiometric reconstructions of human metabolism, such as Recon 2 and Recon 3D, do include amino acid and neurotransmitter metabolism. Recon 2 correctly reproduced the elevated phenylalanine levels in PKU patients [36]. Recon 2 and its successor Recon 3D [37], however, lack the kinetic information that is required to grasp the impact of substrate competition for the LAT1 exchange transporter and the possible existence of other transporters or exchangers. Therefore, to study the consequences of the substrate competition for the LAT1 transporter, we decided to construct a kinetic, ODE-type model.

Here we present the first, detailed, kinetic model of LNAA transport across the BBB, together with the brain dopaminergic and serotonergic metabolic pathways. In this model, we studied the effects of dietary interventions on brain amino acid composition, neurotransmitter metabolism, and protein synthesis in PKU, as related to experimental studies in PKU mice. The model was validated by comparison to a comprehensive dataset from our own research group consisting of plasma and brain amino acid and monoaminergic neurotransmitter levels in PKU mice subjected to several dietary interventions. Furthermore, we simulated the impact of modulating individual dietary amino acids concentrations and the way this could alleviate the pathophysiological cascade towards brain dysfunction as observed in PKU. Since the model is generic, it can be readily applied to other inherited defects of amino acid metabolism and neurological disorders in which the relation between amino acid and neurotransmitter metabolism and the transport across the BBB is important, such as Tyrosinemia type 1 [38], maple syrup urine disease, urea cycle defects, depression [39], autism [40], Alzheimer’s [41], and Parkinson’s disease [42].

## Materials and Methods

### Ethics statement

Experiments were approved by the Ethics Committees for Animal Experiments of the University of Groningen (Permit Number: 6504D).

### Computational methods

The computational model, consisting of a set of 26 Ordinary Differential Equations (ODEs), was built and analysed in Copasi 4.39 [43]. COPASI is a widely used open open-source software package used in modelling biological systems, because it enables easy construction, simulation, and analysis of the models. It has a graphical user interface (GUI) that enables non-expert programmers to study metabolic networks behaviours using many built-in tools for the optimisation, parameter scans, steady state analysis, local sensitivity analysis and metabolic control analysis. Time simulations were performed using the LSODA algorithm for a duration of 100 s simulation time, with relative tolerance of 1·10^-6^, absolute tolerance of 1·10^-12^, and maximally 10,000 internal steps. Steady states were calculated using a combination of methods, according to the default settings in Copasi. The solutions fulfilled the criterion that all-time derivatives of metabolite concentrations approached zero (< 10^-11^). No alternative steady states were found when different initial metabolite concentrations were used. As an input for the steady-state algorithm, the endpoint of the time simulation was used. The detailed model description is available in Text S1. Copasi’s ‘Sensitivities’ algorithm was used to calculate the response coefficients of brain metabolite concentrations to changes in model parameters and blood amino acid concentrations. The model is publicly available together with all the supplementary data in our GitHub repository (https://github.com/WegrzynAB/Papers/ in the folder “2024_mouse_PKU_model_diets”).

### Independent test and validation

As a part of our standard quality control procedures, the model was independently tested by another researcher to assure that the results are reproducible, and the model description (Text S1) agrees to the Copasi script. A thorough comparison between model description and Copasi file was made to check the correctness of all equations and parameter values. Subsequently, a subset of important model simulations was repeated to check if the output reproduced the presented results.

### Dietary interventions in mice

Experiments were performed in BTBR *Pah-enu2* (PKU) and corresponding WT mice, as previously described [19]. In brief, male and female PKU mice received 1 of 5 different LNAA supplemented diets beginning at postnatal day 45. Control groups included PKU mice receiving an isonitrogenic and isocaloric high-protein diet, and PKU and WT mice receiving normal chow. After 6 weeks, brain and plasma amino acid profiles and brain monoaminergic neurotransmitter concentrations were measured.

### Biochemical analysis

Cerebrum and blood samples were processed for the analyses of brain and plasma amino acid and monoamine concentrations, as described previously [27]. Monoamines and related metabolites analysed in the brain included dopamine, norepinephrine, 3-methoxytyramine and normetanephrine in the catecholamine pathway, and serotonin and 5-hydroxyindoleacetic acid (5-HIAA) in the serotonergic pathway.

## Results

### Model construction

We constructed a computational model that describes the transport of LNAA across the BBB and the metabolism of cerebral amino acid and monoamines in mice (Fig.1). The computational model consists of ordinary differential equations (ODEs) and is based on biochemical rate equations. It is focused on the transport of phenylalanine, tyrosine, and tryptophan between blood and brain and on their cerebral metabolism, but also includes the transport of other LNAA to fulfil the requirements for cerebral protein synthesis. We defined three compartments: 1) blood, with fixed amino acid concentrations; 2) the BBB, a cell layer with active protein synthesis, transport processes, and variable amino acid concentrations; and 3) brain, with active protein synthesis [11], transport reactions, monoamine metabolism, and variable metabolite concentrations. Fig.1 gives an overview of all enzymatic reactions and transport processes in the model.

**Figure 1.**
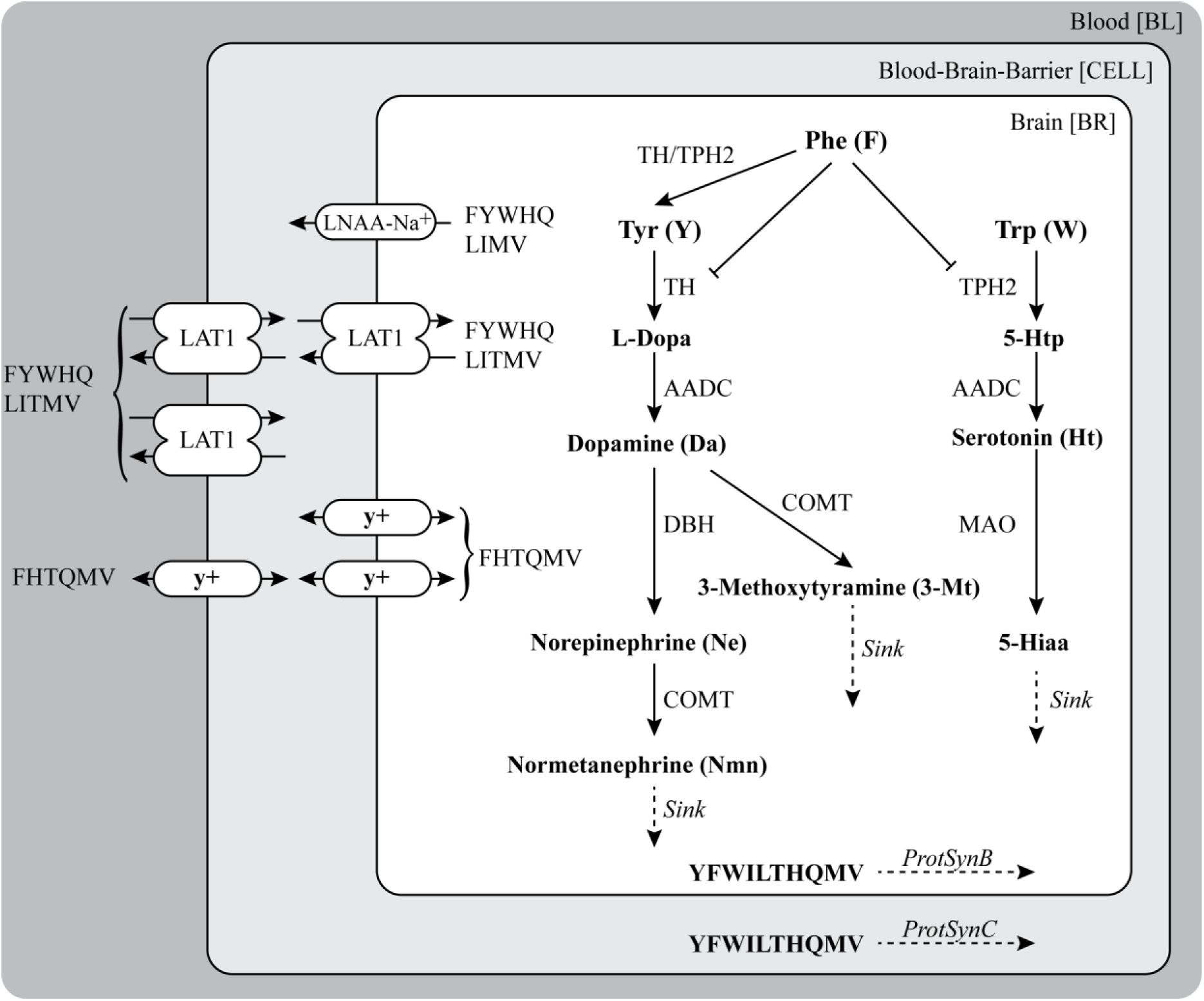
Model of amino acid transport via the blood-brain-barrier and neurotransmitter metabolism inside the brain. The transporter LAT1 is twice as abundant on the luminal (blood-cell) side than on the abluminal (cell-brain) side of the blood-brain-barrier, while the y^+^ transporter displays a reversed distribution. F – phenylalanine, Y – tyrosine, W – tryptophan, H-histidine, Q – glutamine, L – leucine, I -isoleucine, M – methionine, V – valine, T – threonine, 5-Hiaa – 5-Hydroxyindoleacetic acid.

The primary LNAA transporter at the BBB is LAT1. LAT1 is a Na^+^- and pH-independent antiporter, which forms a heterodimeric complex with CD98 glycoprotein and exchanges one amino acid for another. According to Napolitano et al., LAT1 binds the two amino acids on the opposite sides of the membrane in a random order [34]. In the model, the mechanism of LAT1 is described by random-order two-substrates two-products kinetics. This kinetic equation has been extended to account for competition between all LAT1 substrates (Eq. 1 and 2 in Text S1). Since LAT1 is an antiporter, it does not lead to a net import of amino acids. However, due to the different affinities of LAT1 for specific amino acids, it leads to an altered composition of the amino acid pool. In contrast, the so-called y^+^ system is a facilitated diffusion transporter, which catalyses the net transport of LNAA. This transporter shows the highest affinity towards cationic amino acids, but at the same time, it is inhibited by various LNAA [44]. Since no exact mechanism is known, reversible Michaelis-Menten kinetics with an equilibrium constant of 1 and competition between the various substrates was used (Eq. 3 and 4 in Text S1). Finally, we included the Na^+^-dependent large neutral amino acid transporter (Na^+^-LNAA), which actively transports amino acids out of the brain. It transports a range of substrates similar to that of LAT1, but in contrast to LAT1, Na^+^-LNAA is only expressed on the abluminal side of the BBB. Together with the y^+^ system, Na^+^-LNAA controls the total LNAA content of the brain [45]. In our model, the Na^+^-LNAA kinetics are described by an irreversible Michaelis-Menten equation with competition between the various substrates (Eq. 5 in Text S1).

To account for the imbalance of cerebral monoamines, including neurotransmitters, in PKU patients, we included monoamine metabolism in the model (Fig.1). It consists of two main pathways: 1) tyrosine metabolism to L-dopa, dopamine (DA), norepinephrine (NE), 3-methoxytyramine (3-MT), and normetanephrine (NMN); and 2) tryptophan metabolism to 4-hydroxytryptophan (4-HTP), serotonin (HT) and 5-hydroxyindoleacetic acid (5-HIAA). The first step in both pathways is catalysed by an amino acid hydroxylase. Both tyrosine hydroxylase and tryptophan hydroxylase (TH and TPH2, respectively in Fig.1) are inhibited non-competitively and competitively by phenylalanine [46]. Since phenylalanine is not only an inhibitor, but also a substrate for these enzymes, it can be converted at a low rate to tyrosine in the brain [47]. We modified the kinetic mechanism proposed by Ogawa and Ichinose [46] to include not only the role of phenylalanine, but also competitive inhibition by L-dopa [48], NE, and DA [49] for tyrosine hydroxylase, and by 4-HTP [50], L-dopa, and DA [51] for tryptophan hydroxylase (Eq. 6 and 7 in Text S1). Tetrahydrobiopterin was assumed to be available at a saturating concentration for both hydroxylases and therefore not included in the model. Subsequent metabolic steps were described by Michaelis-Menten kinetics, with substrate competition where applicable (see detailed description in Text S1).

Lastly, brain-protein synthesis is affected both in PKU patients [11] and in PAH-deficient mouse models [17]. We included the synthesis of protein starting from the amino acids that were already in the model. Implicitly, we thereby assumed the other amino acids to be present in excess. The amino acid stoichiometry in protein synthesis was calculated from the mouse exome [52]. The affinity constants (K_m_ values) used in the protein-biosynthesis equation reflect the affinity of each of the amino acids to its cognate tRNA-ligase.

All parameters used in the model, except for the Vmax of LAT1, were taken from the literature. Where available, we prioritised murine data, as specified in Table S2 in Text S1. All enzyme rates were normalised per total mouse brain. The value of Vmax of LAT1 enzyme was found by a manual fitting to the WT data, since no value was available that could be related reliably to total mouse protein. Based on the above, we constructed a model of 26 variable metabolite concentrations, 105 reactions, and 89 parameters. Model simulations predicted fluxes and metabolite concentrations both as functions of time and at steady state. Detailed information about the parameter values and their source can be found in the model description (Text S1).

### Experimental validation of the model: the effect of disease and diet on brain amino acids

To better understand the relationship between plasma amino acids and brain biochemistry in PKU, we simulated the effect of different diets that were previously given to PKU mice [53]. The blood concentrations in the model were fixed and set to the values measured in the specific mice groups (see Table S3 in Text S1). PAH, the defective enzyme in PKU, is expressed in the liver and kidney, but not in the brain. The PKU model is therefore distinguished from the WT model by the altered concentrations of amino acids in the blood compartment. Notably, blood phenylalanine is high and blood tyrosine low in PKU compared to WT mice on normal chow diet (Table S4).

Our model distinguishes between the BBB compartment (called CELL) and the brain itself (BR) (Fig.1). The experimental data used for model optimisation and validation, however, had been obtained from total brain samples that also included the BBB. Therefore, we calculated as model output a weighted average of the concentrations in the CELL and BR compartments, considering the difference in the volume of these two compartments (for details see Text S1). Furthermore, since the modelled and experimental data for brain concentrations had different units (µmol/g wet weight and µM, respectively), we compared only values relative to the WT in both experimental data, and model predictions.

The simulated data resembled the experimental data for cerebral amino acid concentrations, based on low normalised difference scores. The most accurate predictions were obtained for glutamine, tyrosine and tryptophan (Fig.S1). The model simulations correctly predicted that cerebral phenylalanine accumulated in PKU mice compared to WT, when both were kept on a non-supplemented chow diet (black bars in Fig.2A). The accumulation of phenylalanine was larger in the model than in the experiment, which is also visible in its large relative difference score (Fig.S1). Nevertheless, the relative increase predicted by the model was within the values reported in the literature (Table 1). Furthermore, we observed a 17% decrease in the protein synthesis rate in the brain compartment and a 27% decrease in the BBB compartment in the PKU model compared to WT, in line with previously published experimental data in PKU mice [17] (Fig.S3).

**Figure 2.**
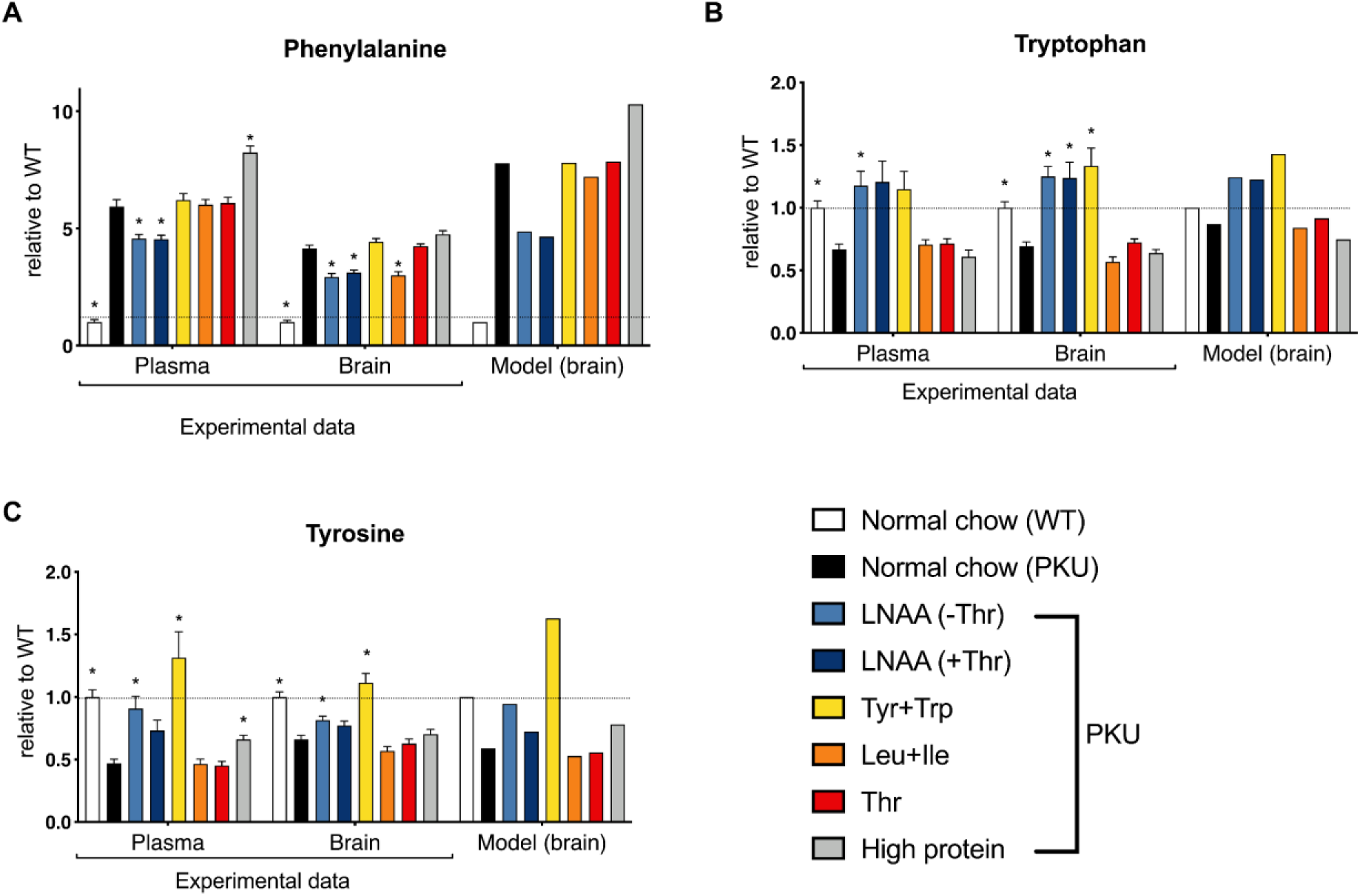
Comparison between experimental data and model predictions for changes in the brain levels of (A) phenylalanine, (B) tryptophan, and (C) tyrosine, in PKU mice with dietary interventions, relative to the WT values. For the experimental data, each bar represents a mean (ns = 16) with a standard error of the man. Significant differences between diets and normal chow PKU mice have been marked with *.

**Table 1.**
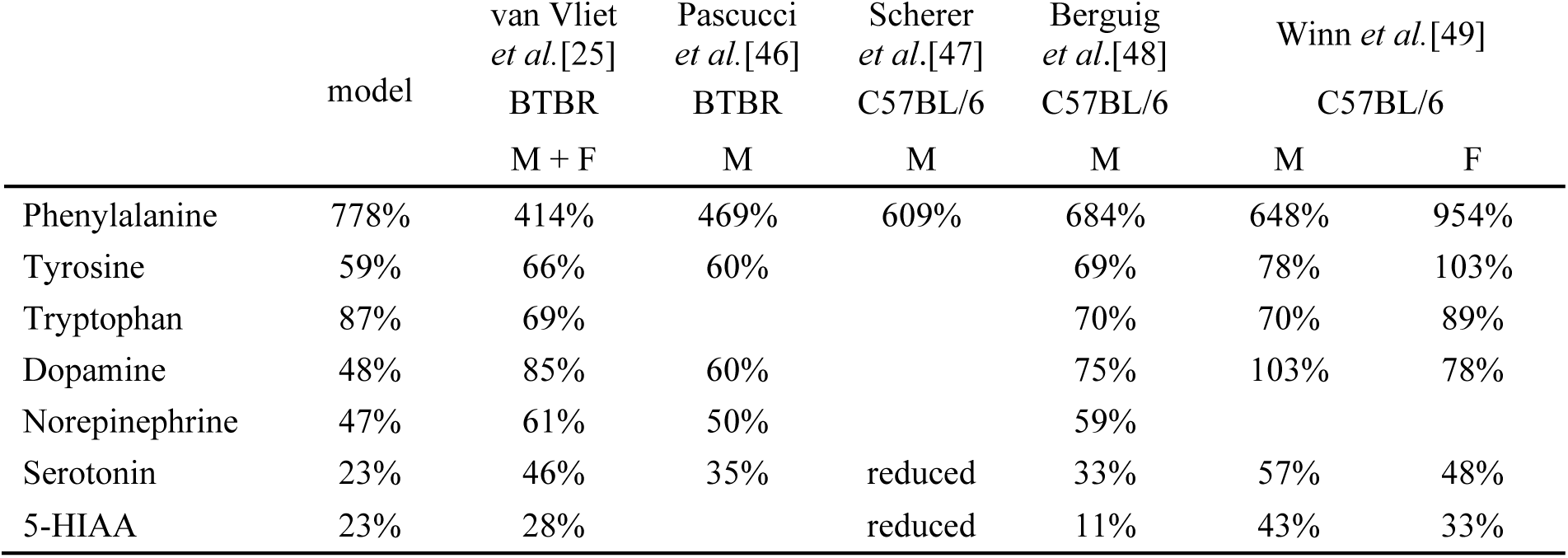
Comparison between the model predictions and literature data on cerebral amino acid and neurotransmitter levels. Relative (PKU/WT) values are shown. BTBR and C57BL/6 denominate the background mice strain, M and F are used to describe males and females, respectively.

Dietary supplementation of PKU mice with all LNAA (with or without threonine) reduced phenylalanine levels in the brain, similarly to what is seen in the experiment (light and dark blue versus black bars in Fig.2A). Furthermore, this intervention restored the protein synthesis rate in the model to the WT levels (Fig.S3). This may suggest that the other LNAAs compete effectively with phenylalanine for transport by LAT1 and other transporters, both in the model and the experiment. We note, however, that the blood concentrations of phenylalanine were also lower in the LNAA supplemented groups (Table S4.4 and [27])). Supplementation of only tyrosine and tryptophan had no effect on brain phenylalanine, neither in the model nor in the experiment (yellow versus black bars in Fig.2A). Experimentally, supplementation of leucine plus isoleucine reduced the phenylalanine concentration in the brain. This effect was attenuated in the model (orange bars, Fig.2A). Previously, the strong impact of leucine and isoleucine was attributed to their high affinities for LAT1 [54]. We have implemented these affinities in the model. In line with this, supplementation of leucine and isoleucine increased their brain concentrations substantially (Fig.S2.). Apparently, however, their high affinities are not sufficient to explain their impact on brain phenylalanine in the experiments, suggesting that leucine and isoleucine act at least in part via another unknown mechanism. Next, threonine supplementation has been given in one study showing a decrease of phenylalanine in blood hypothesising that this decrease in blood phenylalanine levels would also result in a decrease in cerebral phenylalanine [55]. However, this effect was neither seen in experimental data nor in model simulations (red bars, Fig.2A). Finally, a high protein diet served as a positive control in which all amino acids are abundant. Experimental data, as well as model simulations, showed a further increase in the brain phenylalanine levels in PKU mice on a high protein diet (grey bars, Fig.2A).

Subsequently, the impact of the different diets on tryptophan and tyrosine in the brain was accurately predicted by the model (Fig.2B and C). In both model and experiment, tryptophan and tyrosine were most increased by the addition of these amino acids to the diet, while selective supplementation of threonine or leucine plus isoleucine had no effect on the brain concentrations of tryptophan or tyrosine. The brain concentrations of the other amino acids are shown in Fig.S2. First, the concentration of glutamine did not change much on any of the experimental diets in either model or experiment (Fig.S2A). The model missed, however, the increased brain concentration of histidine that was experimentally found in the PKU mice (Fig.S2B). The brain concentrations of leucine and isoleucine did not respond very strongly to either PKU or dietary supplementation of other amino acids. This was qualitatively reproduced by the model (Fig.S2C and D). However, the model predicted a much higher response of brain leucine and isoleucine to supplementation with these amino acids (leucine+isoleucine diet) than the experimental data showed. Finally, the brain concentrations of methionine plus valine (MV) and threonine (Fig.S2E and F) showed similar profiles in the model simulations and the mouse experiment. Particularly, the threonine concentration increased strongly in the brain if supplemented in the diet, either with or without other LNAAs (dark blue and red bars in Fig.S2). Furthermore, except for glutamine and histidine, we observed a correlation between plasma and brain levels of all amino acids, both in the experimental data and in the model predictions (Fig.S2).

### Experimental validation of the model: the effect of disease and diet on neurotransmitters

Subsequently, we compared the brain monoaminergic neurotransmitter levels between model and experiments. The computational model correctly predicted the decrease in both tryptophan- and tyrosine-derived neurotransmitter levels in PKU mice compared to the wild-type controls (Fig.3,4, and Table 1). In general, the decrease was stronger in the model than in the experiments, especially in the dopaminergic pathway (Fig.4, and Table 1). The experimental data shows only a mild decrease in the dopamine levels, followed by a stronger response in the norepinephrine and normetanephrine levels, whilst no change was seen in the 3-methoxytyramine levels. This contrasts with our model predictions where all the dopaminergic metabolites show the same decline in their levels in the PKU model compared with the WT (Fig.4, and Table 1).

**Figure 3.**
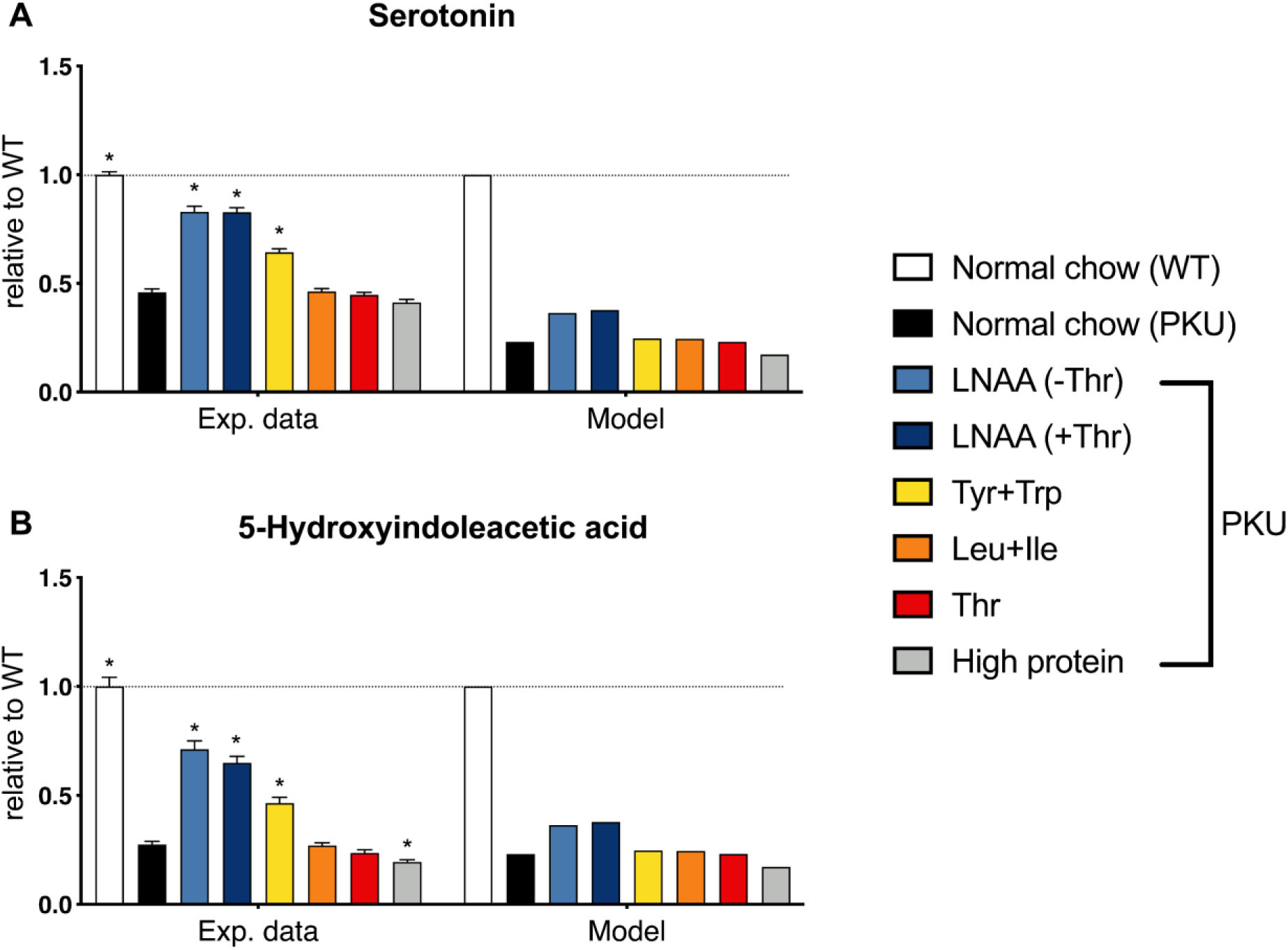
Model validation for the levels of the tryptophan-derived neurotransmitter serotonin (A), and its metabolite 5-HIAA (B) in PKU mice with dietary interventions, relative to the WT values. For the experimental data, each bar represents a mean (ns = 16) with a standard error of the mean. Significant differences between diets and normal chow PKU mice have been marked with *.

**Figure 4.**
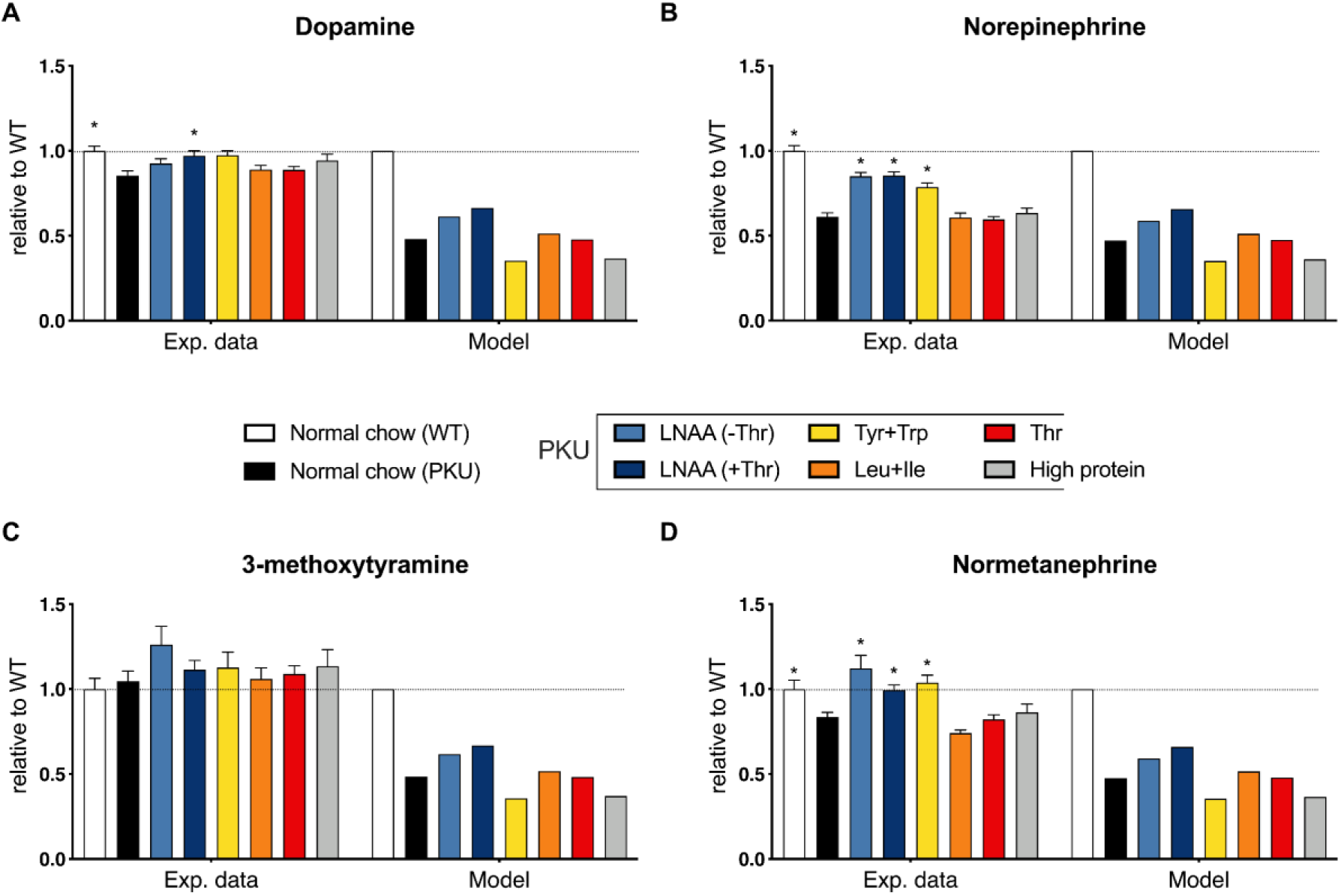
Model validation for the levels of the tyrosine-derived neurotransmitters: (A) dopamine, and (B) norepinephrine, as well as their metabolites: (C) 3-methoxytyramine, and (D) normetanephrine in PKU mice with dietary interventions, relative to the WT values. For the experimental data, each bar represents a mean (ns = 16) with a standard error of the mean. Significant differences between diets and normal chow PKU mice have been marked with *.

Subsequently, we analysed the response of the neurotransmitter concentrations to different diets. Qualitatively, the tryptophan-derived neurotransmitters showed the same dietary profiles in experiments and simulations, but in the simulations the response was attenuated compared to the experiments (Fig.3). In both experiments and simulations serotonin and 5-hydroxyindoleacetic acid levels were effectively increased by supplementation with LNAA plus threonine (Fig.3). In the model, the LNAA diet with threonine increased these neurotransmitters slightly more than the LNAA diet without threonine. The difference was smaller, however, than the experimental error.

According to the model, the LNAA diet with threonine was the only diet with a strong stimulatory effect on the dopamine pathway (Fig. 4). The simulated tyrosine-plus tryptophan-enriched diet even further decreased the levels of the dopaminergic metabolites slightly if compared to the PKU without treatment. This is, however, not seen in the experimental data, which showed a modest increase over normal chow (PKU) in brain dopamine and norepinephrine on the tyrosine-plus tryptophan-enriched diet compared to normal chow. This suggests that we miss a mechanism in the model that buffers the dopaminergic neurotransmitter concentrations in mice *in vivo*.

### Brain amino acid and neurotransmitter levels, and protein synthesis rates are sensitive to the blood concentrations of amino ACIDS

*In vivo*, it is not feasible to assess the impact of each individual amino acid in the blood systematically. Since the model provided a fairly good representation of the *in vivo* situation, we calculated the response coefficients (Fig. 5A) of the clinically relevant output variables towards changes in the concentrations of individual amino acids in the blood. A positive response coefficient means that an increase of the blood amino acid concentration increases the output variable, whereas a negative response coefficient would decrease it.

**Figure 5.**
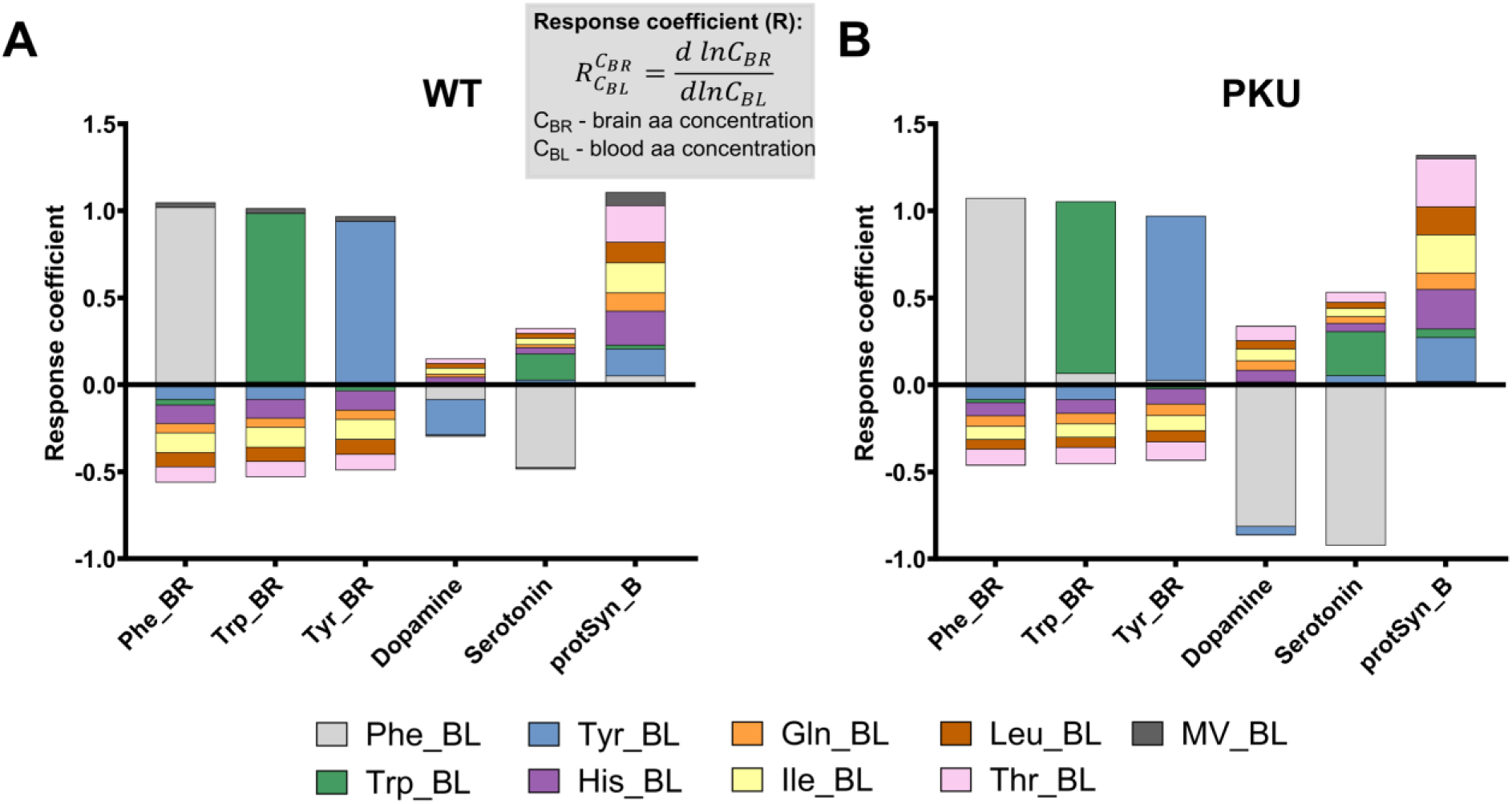
Brain levels of phenylalanine, tryptophan, and tyrosine are susceptible to the changes in the corresponding blood levels of these amino acids in both (A) WT, and (B) PKU models. Bars represent positive and negative response coefficients of brain phenylalanine, tyrosine, tryptophan, as well as dopamine, serotonin, and protein synthesis, to the changes in the blood amino acid concentrations.

The response coefficients were qualitatively similar between WT and PKU model predictions (Fig. 5) as well as in the different diets (Fig. S5). However, in the PKU model, the dopamine and serotonin levels responded more sensitively to the changes in individual amino acid concentrations (cf. Fig. 5A and B). In the following, we will focus on the PKU model (Fig. 5B).

Blood phenylalanine had by far the strongest impact on its own brain concentration as well as on that of the neurotransmitters. The brain tyrosine and tryptophan levels responded only weakly to blood phenylalanine. Surprisingly, these amino acid levels were even increased by increased blood phenylalanine (Fig 5 and Fig. S10). Furthermore, LAT1 reactions showed only a minor concentration control coefficient for brain amino acid levels (Fig. S11). This suggests that competition for LAT1, at least in the present model, is not a dominant factor for these amino acids. The sensitive response of the neurotransmitters, but not of their precursors, suggests that their synthesis is mostly affected by phenylalanine inhibition, rather than by the lack of precursors, at least in the model.

Supplementation of tyrosine or tryptophan had a positive impact on their own brain levels, as indicated by the relatively large, positive response coefficients. Neither had a strong effect on the phenylalanine concentration, confirming that the interaction between phenylalanine on the one hand and tyrosine and tryptophan on the other, was weak. Tryptophan had a positive impact on serotonin, albeit less than the negative effect of phenylalanine. In contrast, the precursor tyrosine had little effect on dopamine. Dopamine is overall less sensitive to blood amino acid levels than serotonin. This may explain why the impact of PKU on dopamine is less than on serotonin in the first place, both in experiments and model. A counterintuitive finding was the negative response of dopamine to plasma tyrosine concentration in the WT, caused by substrate inhibition of tyrosine hydroxylase by tyrosine (Fig. 5B and Fig. S6C). Tyrosine titration showed that this effect was not present in the PKU model since the latter operated at the point where substrate stimulation and substrate inhibition were just balanced (Fig. S6C).

The other amino acids, notably threonine, histidine, leucine, and isoleucine, reduced the brain phenylalanine concentration. Individually they had a low response coefficient, but together they could have a substantial impact (Fig. 5B). They also impacted positively on dopamine and serotonin, presumably through alleviating the inhibition by phenylalanine. To test this hypothesis, we modelled the impact of doubling the plasma concentrations of threonine, histidine, leucine, and isoleucine (THLI perturbation) in the PKU normal chow diet. Furthermore, we tested scenarios where only one of these AAs was increased so that the sum of threonine, histidine, leucine, and isoleucine was the same as in the THLI perturbation. Similarly, we tested the effect of a proportional increase in all non-phenylalanine LNAAs. Last, we tested extension of THLI perturbation by additional supplementation of tyrosine and tryptophan to their WT normal chow plasma levels. As response coefficient analysis suggested, THLI perturbation was able to substantially decrease the brain phenylalanine levels more than any of the amino acids alone (Fig. S12A). However, the effect of THLI perturbation on brain levels of neurotransmitters was only minimal (Fig. S12J-O). Since the leucine only diet showed the least improvement, we decided to test a scenario where only threonine, histidine, and isoleucine (THI perturbation) were added (maintaining the sum of threonine, histidine, and isoleucine equal to the one in THLI perturbation). With THI only supplementation, we saw an even bigger decrease in phenylalanine levels as well as an increase in neurotransmitter levels (Fig S12A and J-O). Additional supplementation of tyrosine and tryptophan in both THLI and THI perturbations showed a further decrease in phenylalanine and an increase in neurotransmitter levels (Fig S12A and J-O).

Furthermore, all amino acids are shown to stimulate protein synthesis in the brain with histidine and isoleucine having the biggest impact in the PKU model (Fig 5B). A drawback of supplementation of these amino acids, however, would be their negative impact on the brain concentrations of tyrosine and tryptophan, most likely through competition for LAT1. However, this may be alleviated by an increase in the tyrosine and tryptophan in the diet (Fig.S12). Interestingly, histidine-only perturbation showed that at high brain histidine concentrations the weak inhibition of DBH enzyme by histidine could lead to a reduction of norepinephrine and normetanephrine levels while dopamine levels remain stable Fig. S12J-M).

Increase in the blood phenylalanine levels alone does not decrease the brain tyrosine and tryptophan levels, but it does inhibit the synthesis of neurotransmitters.

In PKU mice, the blood concentrations of multiple amino acids were changed simultaneously compared to the WT. To disentangle the effect of the increased phenylalanine concentration from that of the other amino acids, we performed an *in silico* experiment. To this end, all blood amino acid concentrations were fixed at the levels measured in either WT or PKU mice without dietary supplementation. Then, Phe was increased to PKU level in the WT (Phe 1803, corresponding to 1803 μM), or to WT level in the PKU model (Phe 304, corresponding to 304 μM) (Fig. 6). Brain phenylalanine accumulates in response to the change in the blood phenylalanine levels as expected (cf. WT Phe 304 to WT 1803; or PKU Phe 304 to PKU 1803 in Fig. 6A). In contrast, brain phenylalanine did not respond to other amino acids that were altered in PKU (cf. WT Phe 304 to PKU Phe 304; or WT Phe 1803 to PKU Phe 1803 in Fig. 6A). Tyrosine and tryptophan hardly responded to the change in phenylalanine but decreased in response to alterations of the other amino acids in PKU (cf. WT to PKU at either Phe concentration in Fig. 6A).

**Figure 6.**
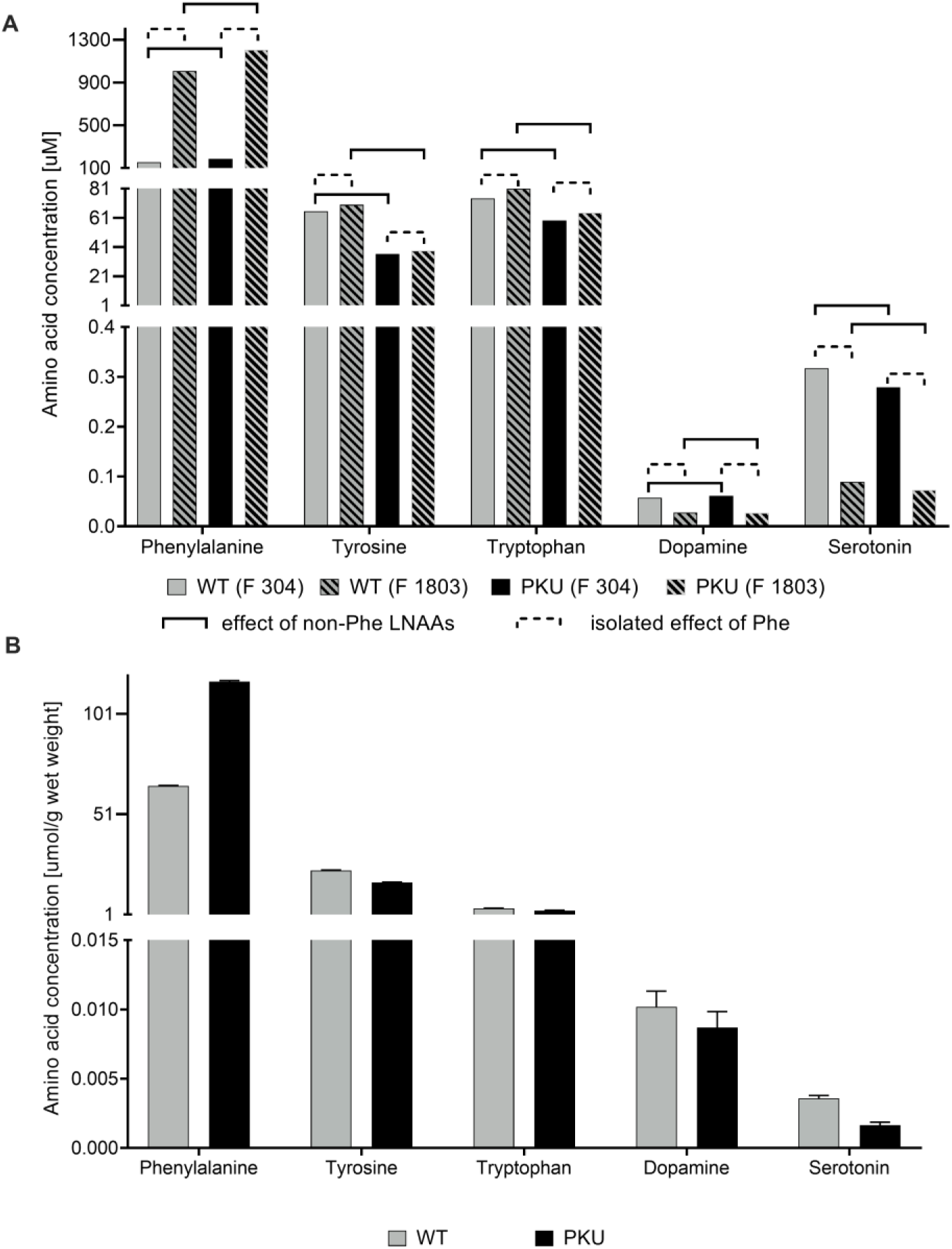
Changes to the brain phenylalanine, tyrosine, tryptophan, dopamine, and serotonin levels in response to the blood phenylalanine increase/decrease in WT and PKU model simulations (A), in comparison with the experimental data from mice (B). WT or PKU indicate that all blood amino acid concentrations except those of phenylalanine were equal to those measured in WT or PKU, respectively. Phe 304 indicates that the WT phenylalanine concentration of 304 μM was used, whereas Phe 1803 indicates that the PKU phenylalanine concentration of 1803 μM was used.

The brain neurotransmitters decreased strongly in response to the increased blood phenylalanine levels (Fig. 6A and Fig. S7C and D) in line with the experimental data (Fig. 6B). However, the dopamine to serotonin ratio in the model was opposite to that of the one seen in the experimental data (Fig. 6B). Even when the blood concentrations of tyrosine and tryptophan were doubled or quadrupled, the strong inhibition of dopamine and serotonin levels by phenylalanine synthesis persisted (Fig. S7-S9). Only when blood phenylalanine concentration was lowered to 750 μM (Phe 750, halfway between WT and PKU values) the combined addition of tyrosine and tryptophan had a positive effect on serotonin, while dopamine levels were still lower than in the WT (Phe 750, Fig. S9).

## Discussion

In this work, we present the first dynamic model that links the LNAA transport system across the BBB to the subsequent brain neurotransmitter metabolism and protein synthesis (Fig. 1). In particular, we included extensive competition between the substrates of the LAT1 transporter across the BBB. The model was designed to ultimately understand and optimise how dietary treatment affects the brain biochemistry in PKU patients.

As the mouse brain is experimentally more accessible than the human brain, and PKU mouse models closely resemble the genetics, biochemistry, and neurobiology of human PKU, the detailed mechanism of PKU pathophysiology has mostly been studied in PKU mice. Therefore, the model was based on data from the BTBR mouse strain and validated against data of the PKU mice in the same background strain. Considering the large number and heterogeneity of the kinetic parameters (Table S2) the correspondence between model predictions and experimental data was remarkably good. Moreover, while not all metabolic fates of Phe, tyrosine and tryptophan were included in the model, we included the most important metabolic pathways. Kinetic parameters were not fitted to the desired outcome but based on biochemical data of specific enzymes. Models of this type give insight in how complete our biochemical knowledge is to understand functional properties, such as the neurotransmitter levels [56,57].

Qualitatively, the model largely reproduced the impact of PKU and most of the dietary treatments on brain biochemistry (Figures 2-4, and Table 1) as seen in the mice with PKU. The most conspicuous exception was the leucine plus isoleucine diet, which reduced phenylalanine levels in PKU mouse experiments, but barely in the PKU model. The affinity of LAT1 for leucine and isoleucine in the model was high, in accordance with the biochemical data, and these amino acids readily crossed the BBB. Yet, their calculated impact on phenylalanine uptake was limited. This discrepancy led us to conclude that leucine and isoleucine probably affect brain phenylalanine levels via another mechanism besides the simple competition for LAT1. So far, existing research on the metabolic role of branched-chain amino acids shows that leucine can stimulate protein synthesis and decrease protein breakdown [58,59]. Indeed, in our simulations, we saw a small increase in protein synthesis rate in leucine and isoleucine diet (Fig. S3). Since phenylalanine is more abundant than tyrosine and tryptophan in mouse protein [52], stimulation of protein synthesis could decrease free phenylalanine levels, without depleting the free tyrosine and tryptophan levels. Another possibility might be that LAT1 is not the only transporter but that other transporters - such as LAT2 of which the function and the location of in brain is still debated - are not sufficiently considered.

Another qualitative discrepancy between model and experiment was the finding that the tyrosine plus tryptophan diet increased the neurotransmitter levels experimentally, but not in the model. This might well be explained by the fact that, in the model, brain phenylalanine levels in PKU compared to WT were much more increased than seen in the PKU mouse experiments. On the contrary, the reduction of brain tyrosine and tryptophan levels in PKU compared to WT in the model was less than observed in the mouse experiments.

Quantitatively, the calculated effect of PKU on most brain metabolites in the model was larger than the measured effects in the mouse experiments, with was especially true for brain phenylalanine levels. The fundamental reason why a quantitative agreement is beyond reach at this stage could be the compartmentation of the brain. The available validation data are in µmol/g wet weight of the total brain, whereas the model predicts local concentrations in µmol/L. The model already includes some compartmentation, specifically the blood compartment, the endothelial cells of the BBB, and the brain itself. To relate the model outcome to the validation dataset, a weighted average between the BBB and the brain compartment was made and an estimate of the cytosolic volume relative to the brain wet weight. Most likely, the uncertainty in this conversion is an important reason for the quantitative discrepancy. Naturally, uncertainties in the biochemical parameters may also play a role, taken into account that only few of the parameters were mouse specific. However, not all the parameters exert strong control on the brain concentrations of neurotransmitters and amino acids, and in general, we found the biochemical literature of high quality. Further progress could be made by using an intermediate experimental system that is closer to the model. Taslimifar et al. [30,35] proposed further compartmentation of their brain model into blood, endothelial cells, cerebral spinal fluid, and dopaminergic and serotonergic neurons. To validate such a model experimentally, novel organ-on-chip technology holds great promise, particularly since different tissue- and or cell-type-specific chips can be coupled functionally [60,61].

At this stage, the qualitative agreement between model and mice experiments already allows us to interrogate the model. For many mechanistic and clinical questions, it is sufficient to rank the impact of different interventions and to predict in which direction they could work. Given the complexity of the system, modelling can provide non-trivial answers. A particular advantage is that, in a model, the impact of individual amino acids in the blood can be investigated one by one, in contrast to the different diets which change multiple amino acid levels in the blood at the same time.

Firstly, the model correctly predicted the altered brain levels of amino acids in PKU (Fig. 2). Notably, the phenylalanine concentration was increased, whereas tyrosine and tryptophan were decreased in the brain. The model suggests that the decrease of cerebral tyrosine and tryptophan is not primarily due to competition with the high phenylalanine concentration for the LAT1 transporter. Rather, tyrosine and tryptophan were already decreased in the blood (Table S4). According to the model, this decrease of blood tyrosine and tryptophan appeared to be a prerequisite for the decrease of their cerebral concentrations (Fig. 6). The lower blood levels of tyrosine in PKU mice and patients can be attributed to impaired production in the liver due to the PAH deficiency, but may also be caused by alterations the intestinal uptake, altered microbiota composition or metabolism [62,63]. The lower blood levels of tryptophan in PKU mice and patients are less well understood, but altered metabolism of tryptophan via the kynurenine pathway has been suggested to play a role [64].

Second, the model qualitatively reproduced the decline of brain neurotransmitter levels that was observed in PKU mice [19,65]. The fact that the dopamine pattern mimics that of tyrosine, while the serotonin pattern mimics that of tryptophan, may seem to suggest that neurotransmitter levels are controlled by precursor levels. However, the response coefficients in the model (Fig. 5) showed the opposite: both cerebral serotonin and dopamine were strongly negatively controlled by blood phenylalanine, most likely via the strong negative inhibition of tyrosine and tryptophan hydroxylases by phenylalanine (Fig. 1).

Third, the response coefficients showed that isoleucine, leucine, histidine, threonine, and tyrosine all contributed individually to a reduction of the phenylalanine levels and an increase of neurotransmitter levels in the brain of PKU mice (Fig. 5, Fig. S12). This is surprising since neither the leucine plus isoleucine nor the threonine diet affected brain phenylalanine and neurotransmitter levels in the model (Fig. 2). We should keep in mind, however, that the effect of individual amino acids that compete for LAT1 was very small compared to that of phenylalanine. When supplemented together, however, as in the complete LNAA supplementation, they have a strong impact (Fig. 2 and 3, Fig. S12).

A final striking result was the weak, but significantly positive, effect of blood phenylalanine on the brain concentrations of tyrosine and tryptophan in the model. This was seen in the positive response coefficients of blood phenylalanine on tryptophan in PKU (Figure 5B) as well as in a minor increase of tryptophan and tyrosine when the impact of elevated phenylalanine in PKU was simulated without the concomitant decrease of the other amino acids (Figure 6A). The effect can be explained from the fact that LAT1 is an antiporter; phenylalanine does not only compete for LAT1 at the blood side but also serves as a counter metabolite at the brain side as seen in the increased rates of tyrosine and tryptophan transport to the brain with an increase of phenylalanine (Figure S10). The inverse effect was not observed: when tryptophan or tyrosine were increased separately in the model, their negative effect through competition for LAT1 dominated the uptake of phenylalanine (Figure 5B).

What do these results mean for a clinical application of the LNAA diet? To answer this question, we must emphasise that we did not simulate the altered diets used in the PKU mice per se, but rather the impact of altered blood concentrations of amino acids on the brain in the PKU mice. For a more complete insight into the impact of diets, we should also include intestinal uptake and passage through the liver. Nevertheless, the model may help to pinpoint blood amino acids that are important to monitor and optimise in LNAA treatment.

Our present modelling confirmed that the clinical importance of reduction of blood phenylalanine is not only to avoid direct phenylalanine toxicity in the brain but also - albeit to an unknown degree of importance - to reduce the inhibition of tyrosine and tryptophan hydroxylases by cerebral phenylalanine. Furthermore, we gave a theoretical underpinning of the LNAA diet: even though each individual amino acid had a small effect on brain phenylalanine, together they had a strong impact. On the other hand, some discrepancies have been identified between our previous experimental data and the current modelling. These discrepancies reveal the gaps in our current understanding of the pathophysiological mechanisms underlying brain amino acid and neurotransmitter deficiencies in PKU. Our work suggests that future studies should focus on the mechanism through which the leucine plus isoleucine diet reduces brain phenylalanine, on brain compartmentation, and on the kinetic regulation of neurotransmitter biosynthesis. Increased future understanding of these mechanisms should aim to provide an even more solid advice on the optimal LNAA content for trials in PKU patients.

In conclusion, this first detailed, dynamic model of the LNAA transport and subsequent brain neurotransmitter metabolism gives a good, albeit qualitative, description of the impact of dietary treatment of PKU mice. In the future, it may be optimised towards the human patient situation. This can be readily done by changing the model parameters to be human-specific, since the biochemical architecture of the network is thought to be the comparable between mice and man [66]. Moreover, its generic nature makes it applicable to other diseases in which the balance of amino acids and neurotransmitters is affected, such as Alzheimer’s [41] or Parkinson’s Disease [42], depression [39], and autism [40].

## SUPPLEMENTARY INFORMATION

**Figure S1.**
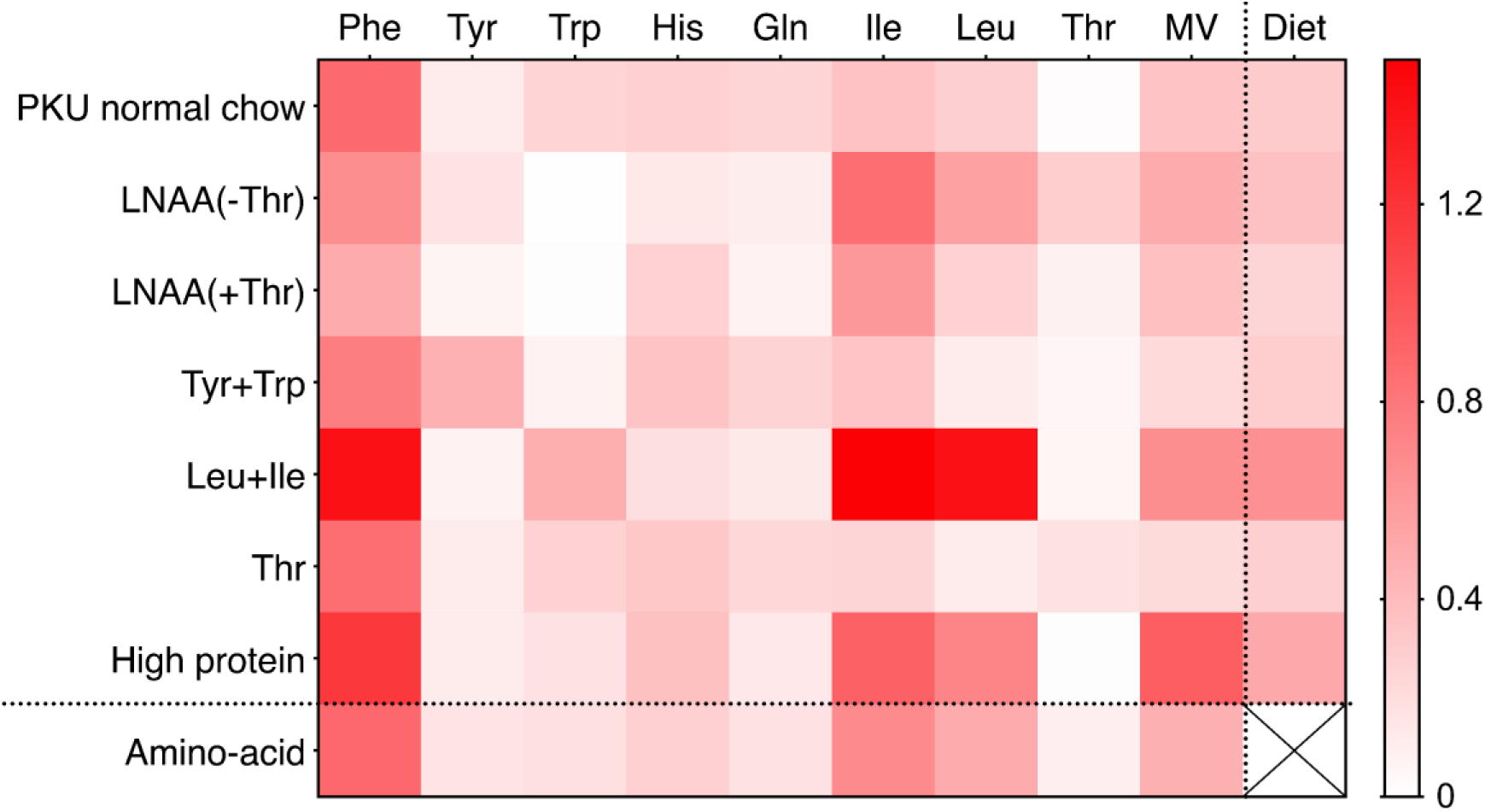
A detailed comparison between model prediction and experimental data. Heatmap shows normalised difference scores between the model prediction for a specific amino acid value relative to WT, and its experimental value relative to WT, for each diet. *Normalised difference score_n,n_* = (|*norm.simulation_n,m_ – norm.data_n,m_*|) · *norm.data*^−1^_*n,m*_, where n = specific amino acid, m = specific diet. If normalised difference score = 0 data and simulation are identical.

**Figure S2.**
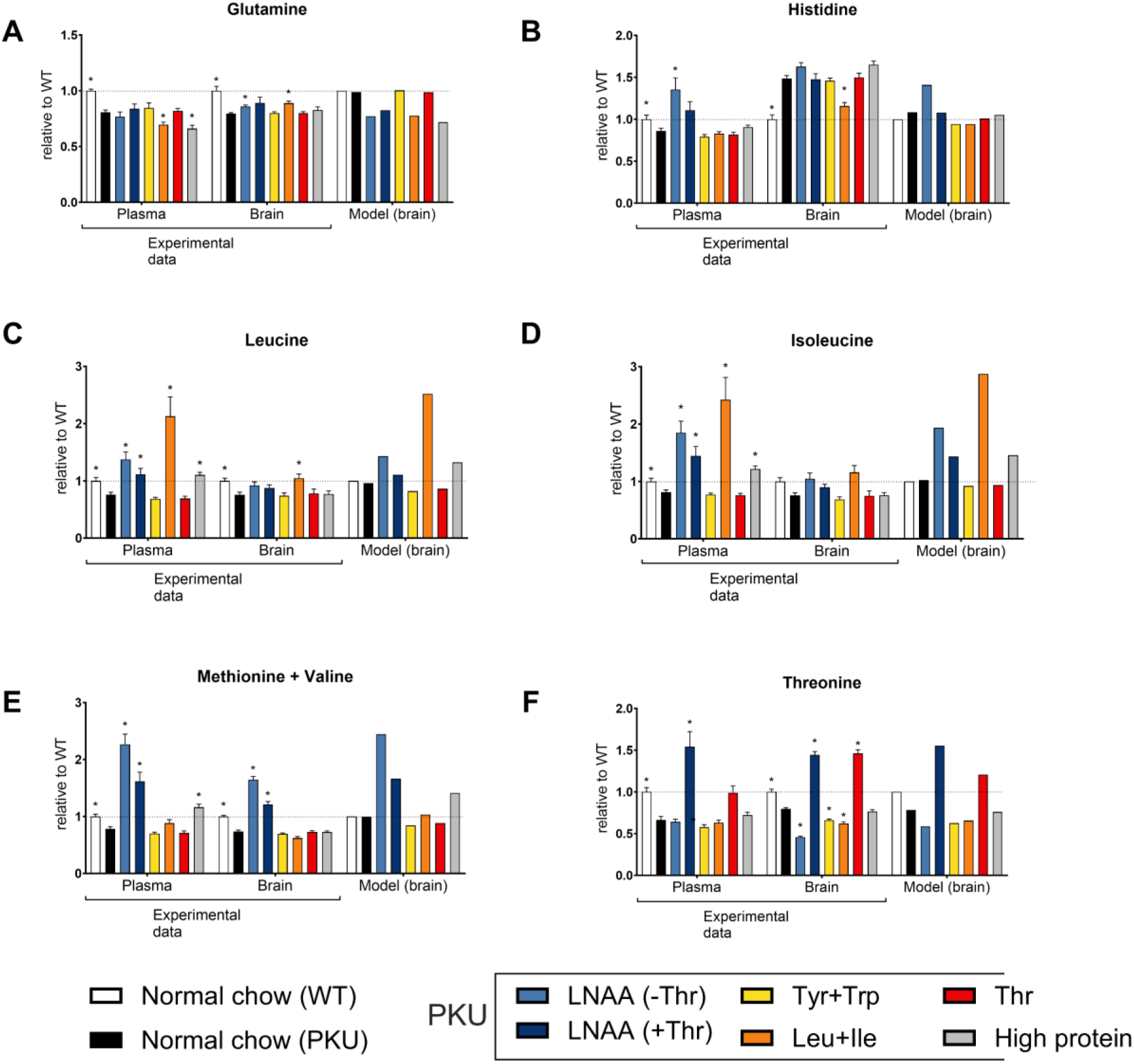
Comparison between experimental data and model predictions for changes in other amino acid levels after the dietary intervention. For the experimental data, each bar represents a mean (ns = 16) with a standard error of the mean.

**Figure S3.**
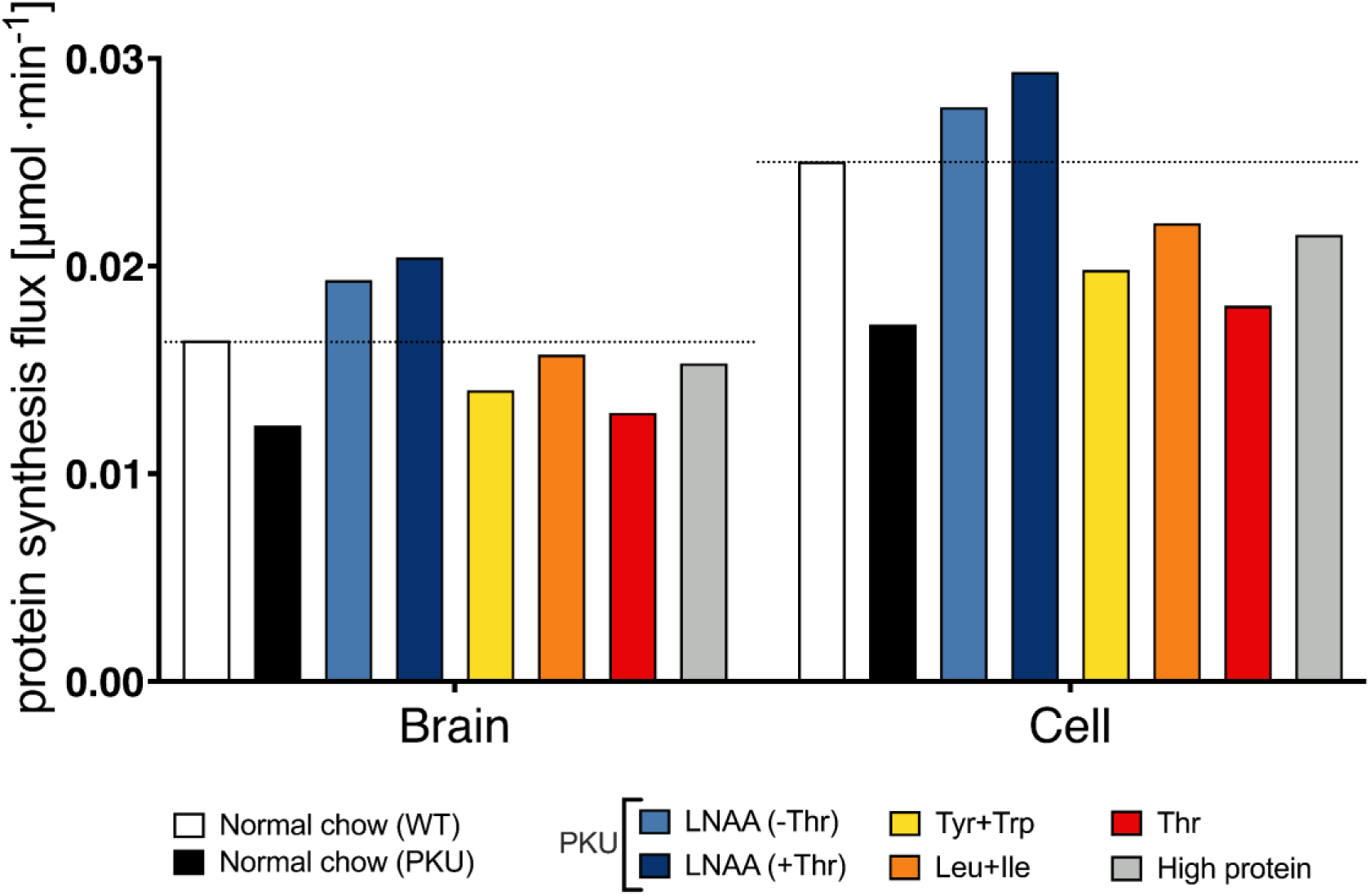
Comparison between the protein synthesis flux in the model in response to different diets.

**Figure S4.**
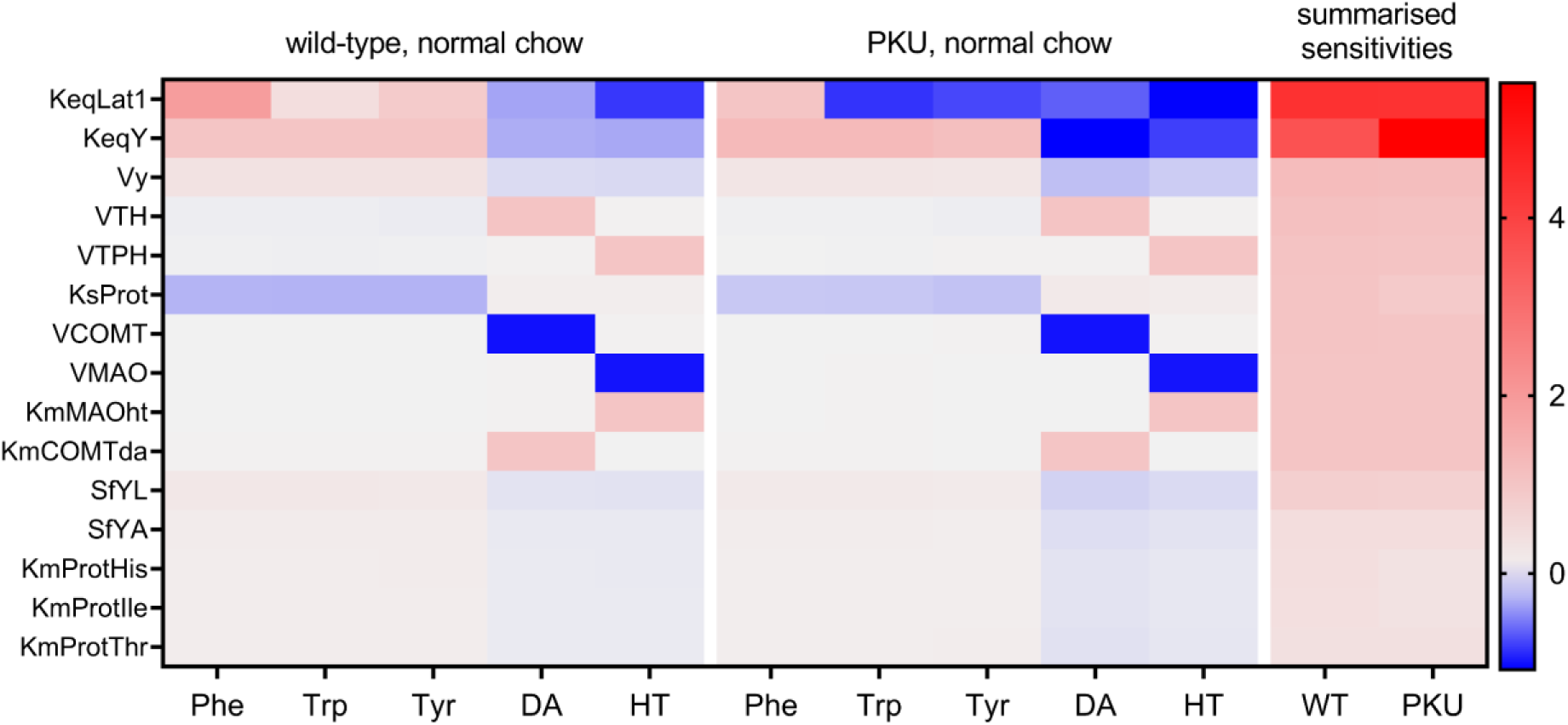
Sensitivities of the brain amino acids to the changes in the model parameters. Top 15 parameters with the most control over the amino acids and neurotransmitters concentrations in WT mice are shown.

**Figure S5.**
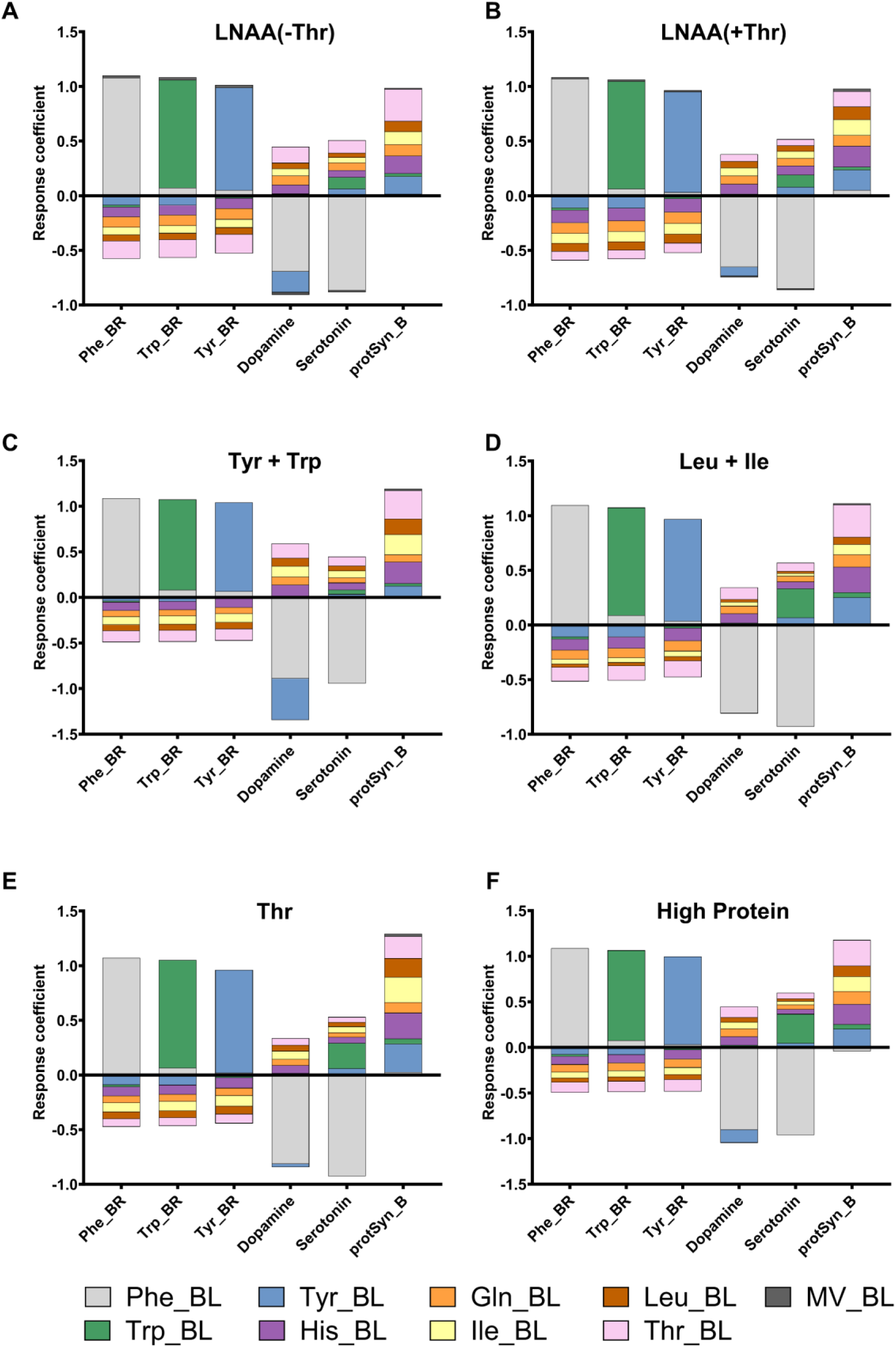
Brain levels of phenylalanine, tryptophan, and tyrosine are susceptible to the changes in the corresponding blood levels of these amino acids in other diets. Bars represent positive and negative response coefficients of brain Phe, Tyr, Trp, as well as dopamine, serotonin, and protein synthesis in the brain (protSyn_B), to the changes in the blood amino acid concentrations. Each graph represents response coefficients calculated based on different dietary conditions as starting points.

**Figure S6.**
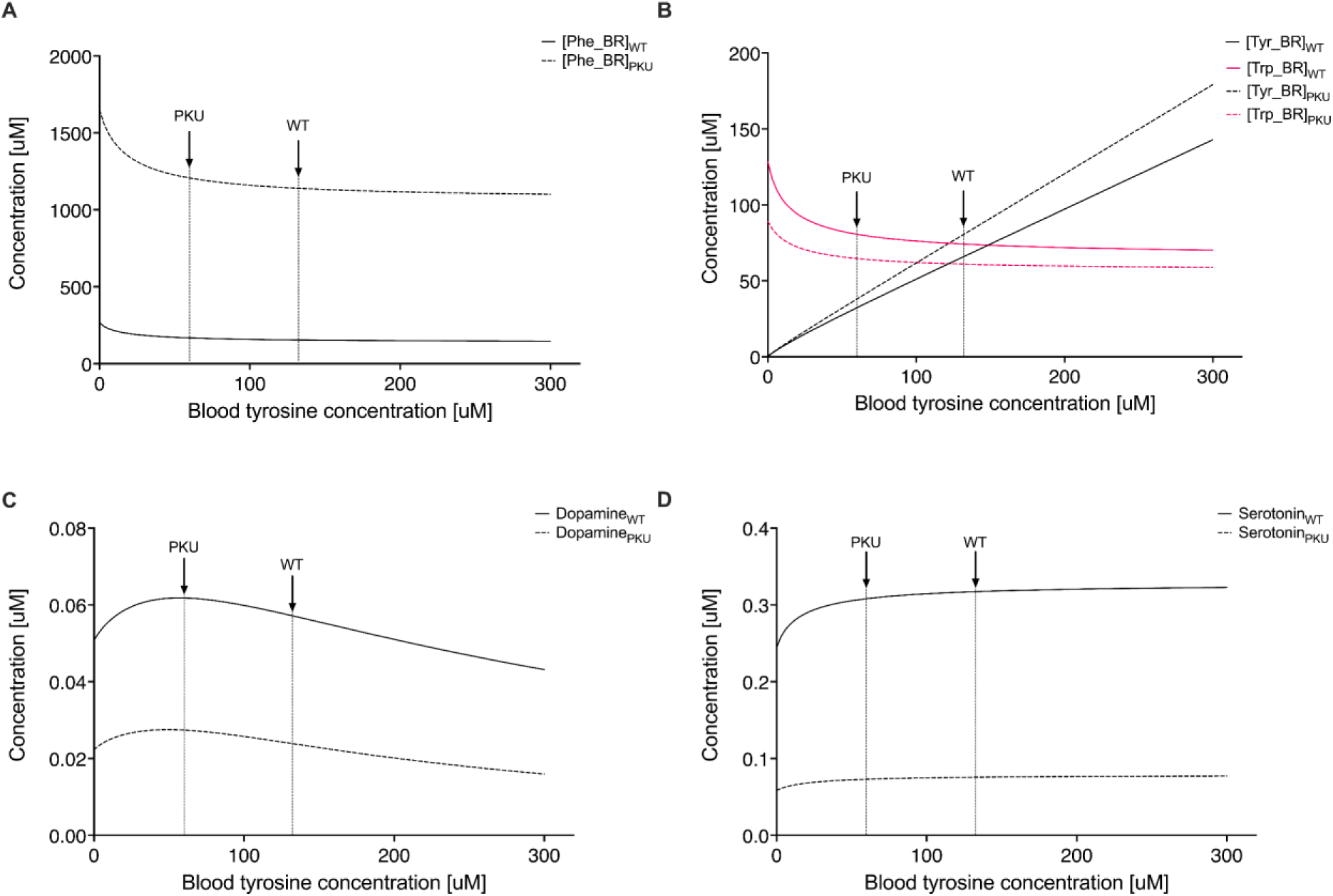
Changes to the amino acid and neurotransmitters concentrations in response to the increasing concentration of tyrosine in the brain. All other amino acid concentrations were fixed at the levels measured in WT or untreated PKU mice, respectively. The arrows indicate the blood concentrations of tyrosine and tryptophan in WT and PKU mice.

**Figure S7.**
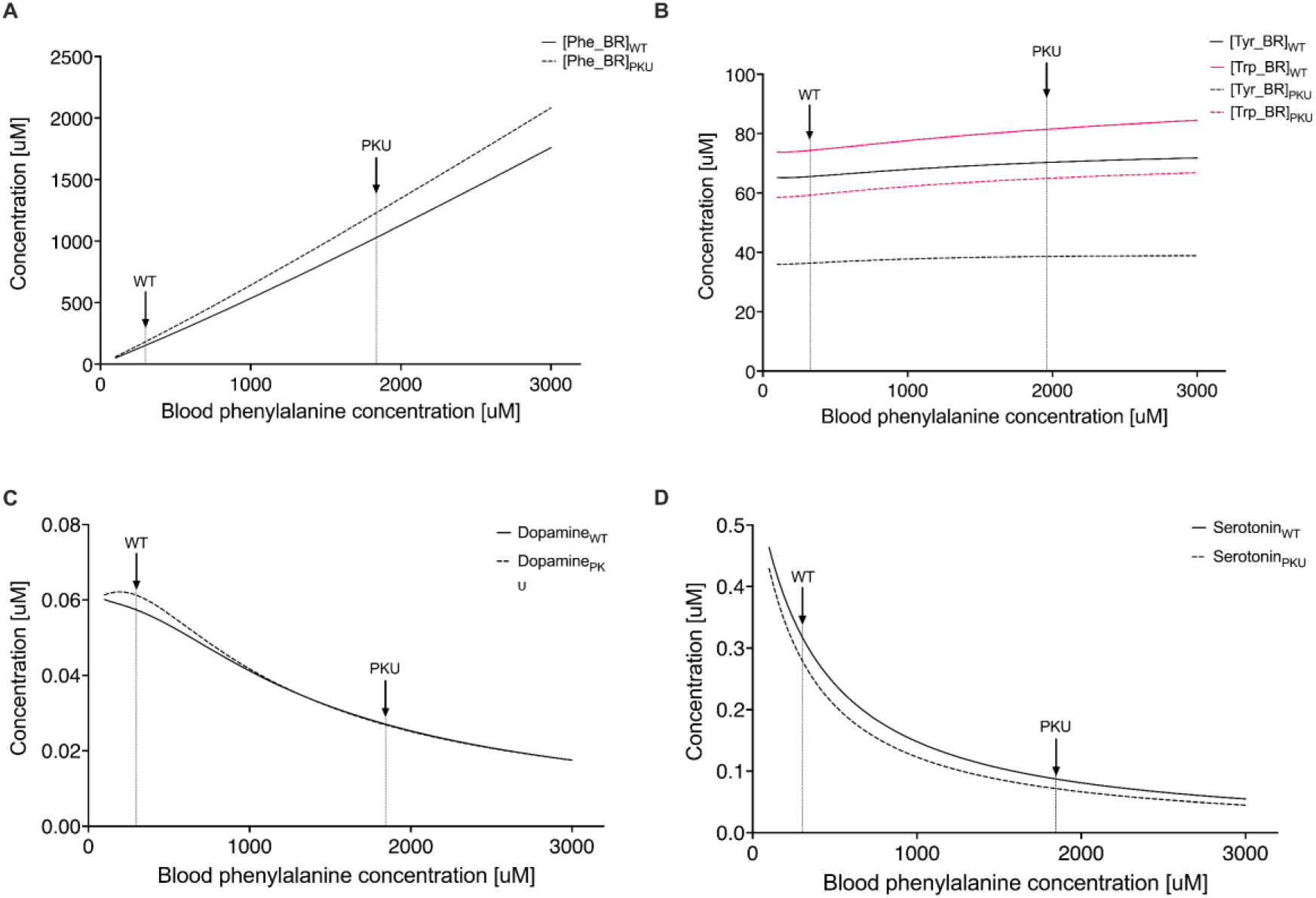
Changes to the amino acid and neurotransmitters concentrations in response to the increasing concentration of phenylalanine in the brain. All other amino acid concentrations were fixed at the levels measured in WT or PKU mice, respectively. The arrows indicate the blood concentrations of tyrosine and tryptophan in WT and PKU mice.

**Figure S8.**
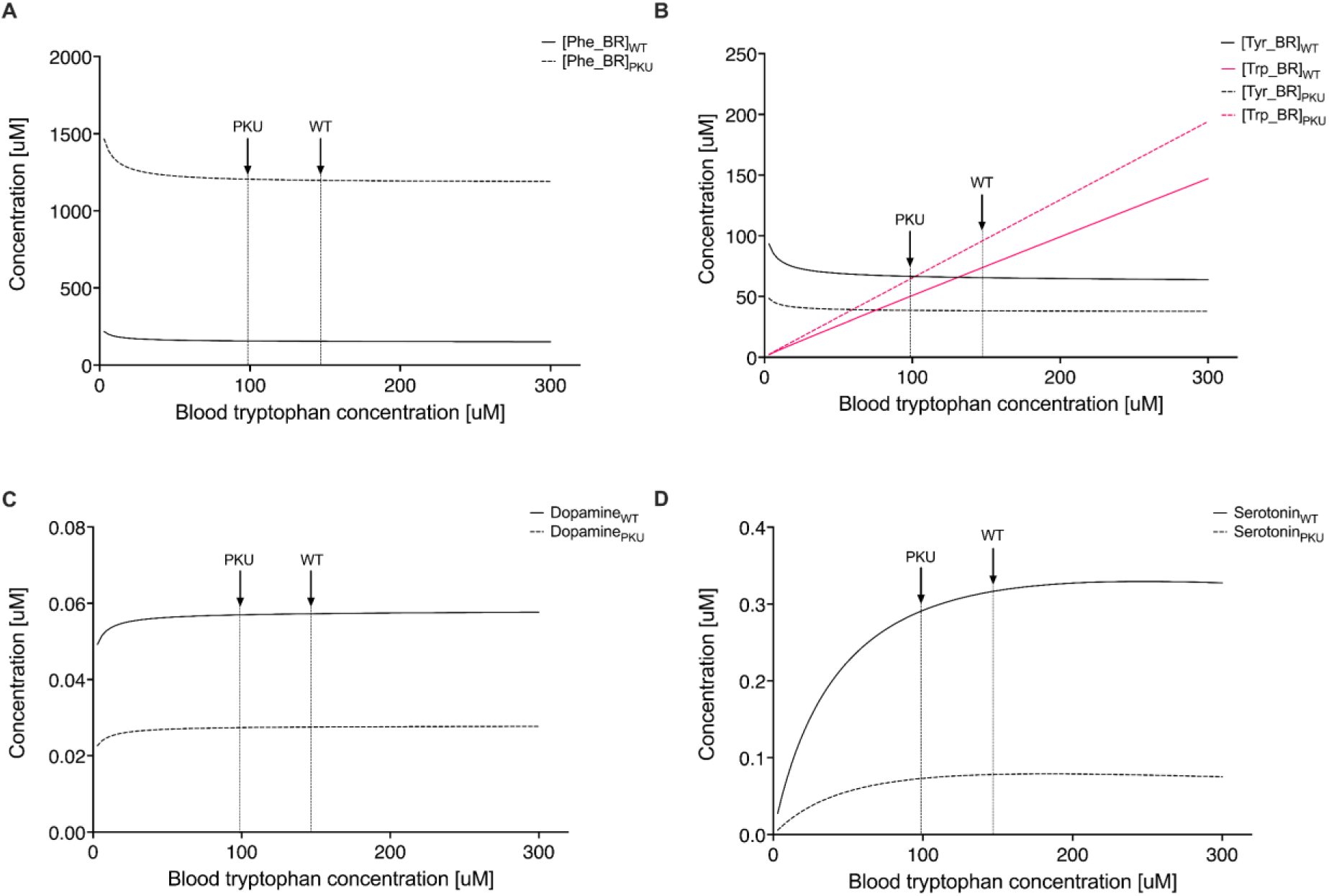
Changes to the amino acid and neurotransmitters concentrations in response to the increasing concentration of tryptophan in the brain. All other amino acid concentrations were fixed at the levels measured in WT or PKU mice, respectively. The arrows indicate the blood concentrations of tyrosine and tryptophan in WT and PKU mice.

**Figure S9.**
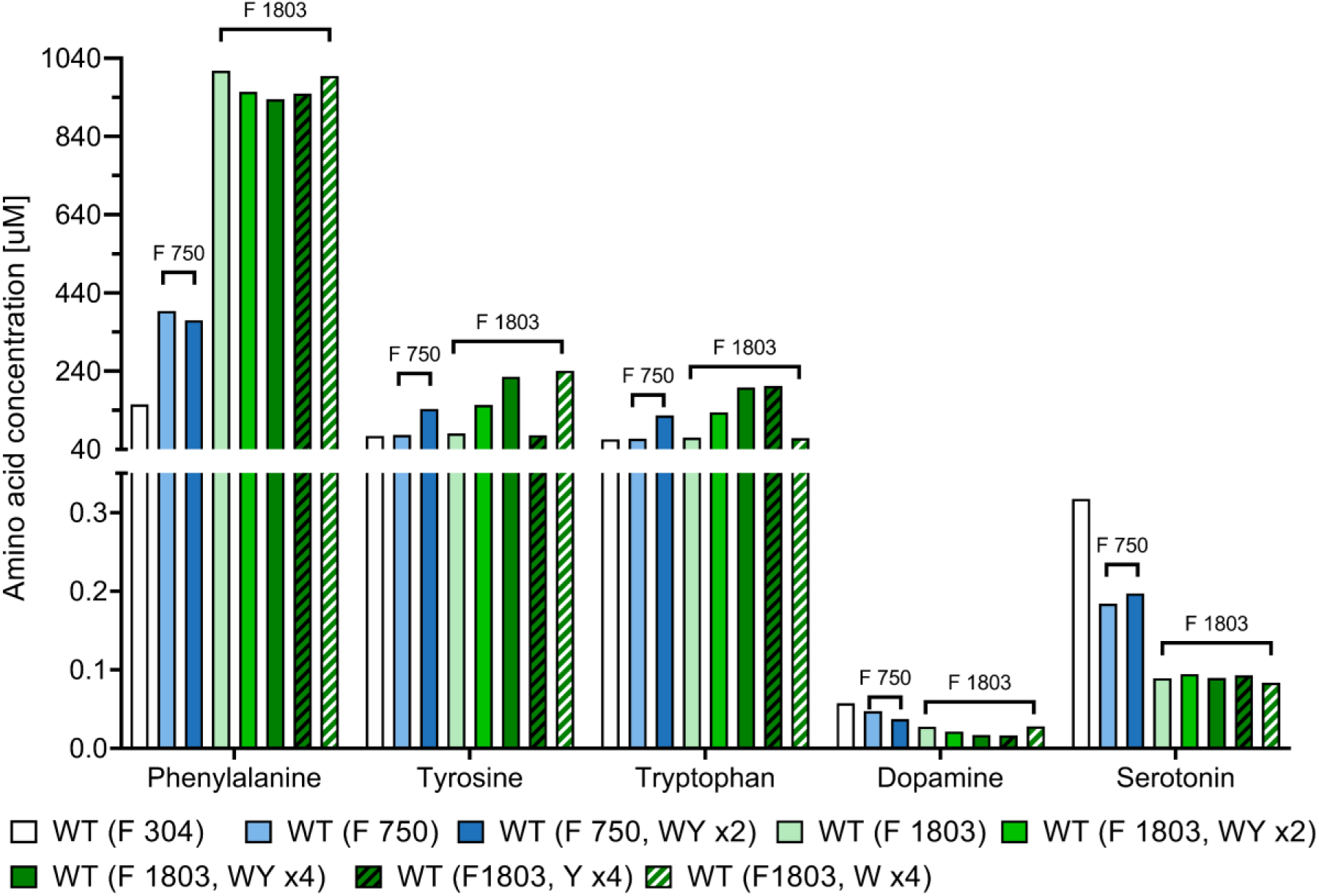
Changes to the amino acid and neurotransmitters concentrations in response to phenylalanine (F) increase and addition of tyrosine (Y) and tryptophan (W) in the WT background.

**Figure S10.**
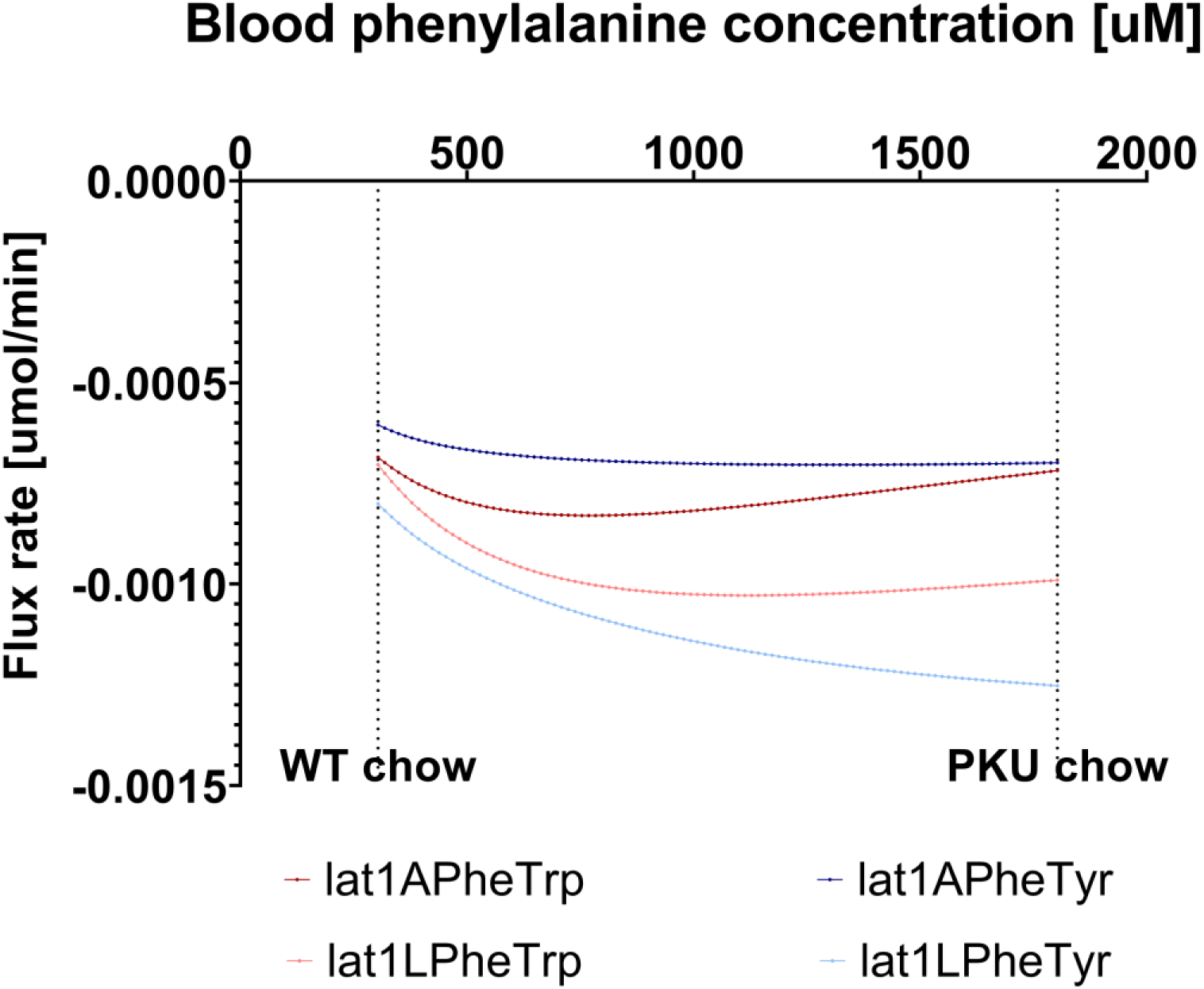
Flux rates of LAT1 exchange between phenylalanine and tryptophan and tyrosine at increasing blood phenylalanine concentrations in WT background.

**Figure S11.**
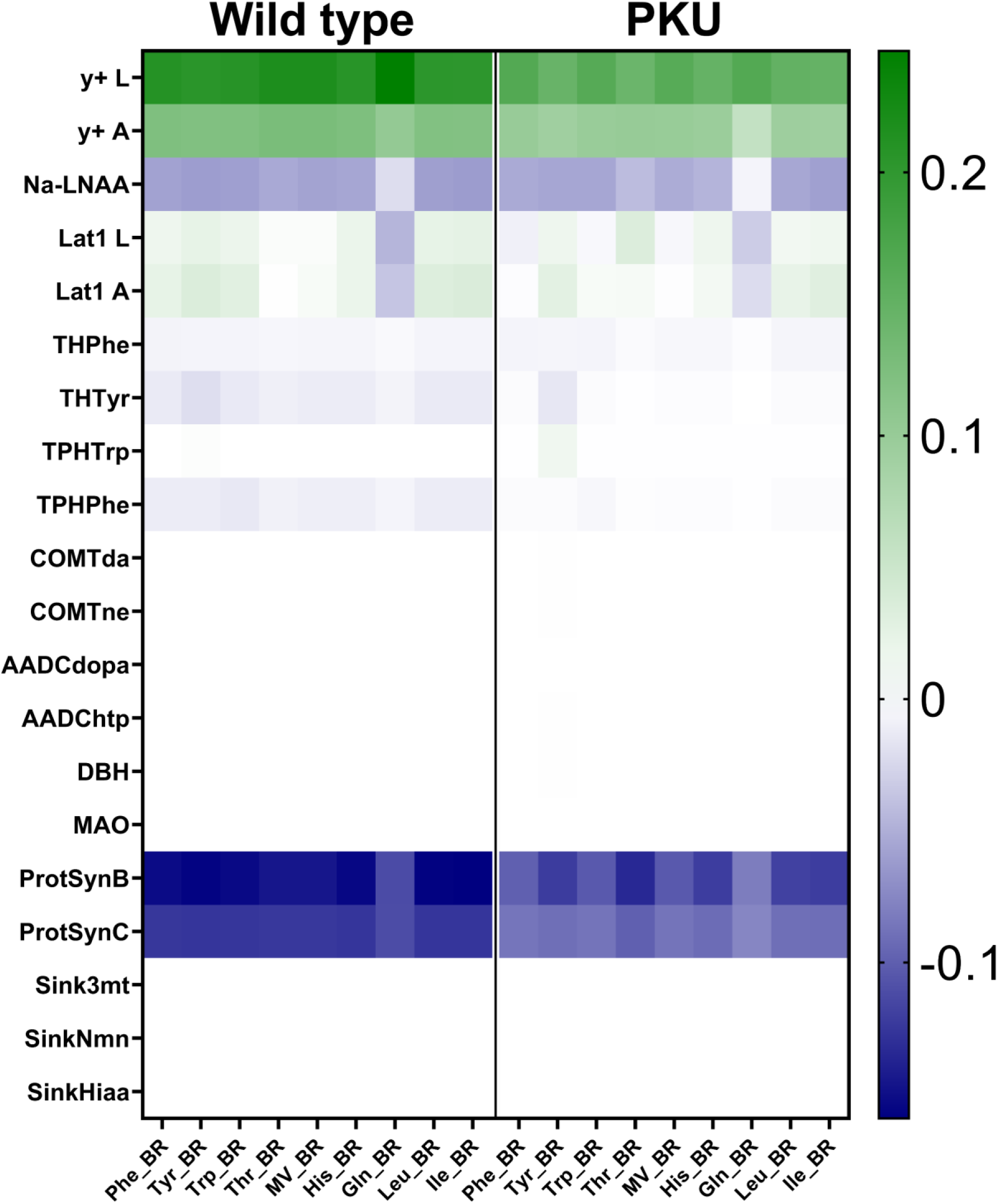
Heatmap of concentration control coefficients for all reactions at WT normal chow AA plasma concentrations, and PKU normal chow AA plasma concentrations. Concentration control coefficients for reactions of transporters have been summarised.

**Figure S12.**
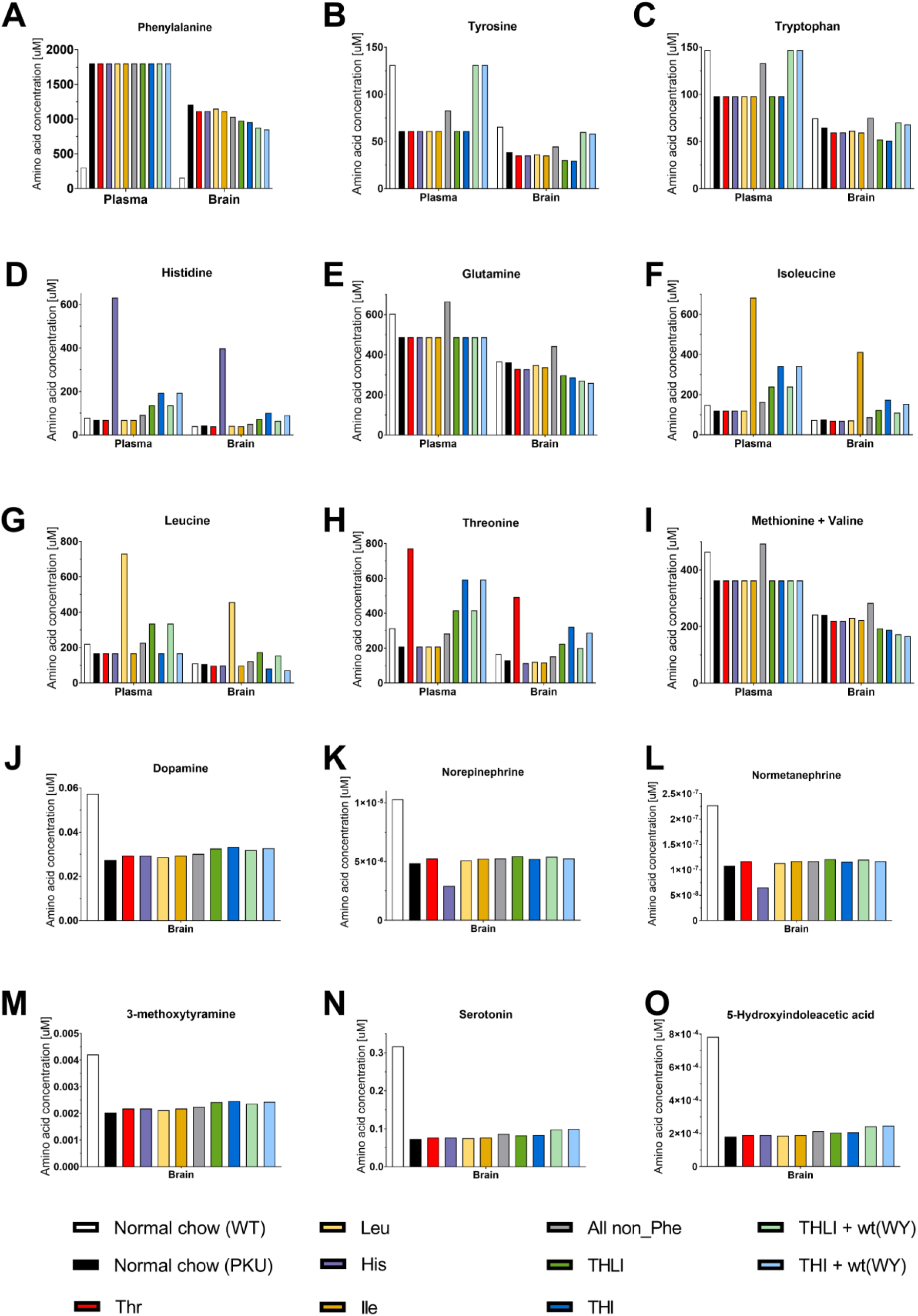
Changes to the amino acid concentrations in response to changes in the plasma amino acid composition. All non_Phe – non-Phe LNAA; THLI – threonine, histidine, leucine, isoleucine; THI -threonine, histidine, isoleucine; THLI + wt(WY) – THLI with wild type levels of tyrosine and tryptophan; THI + wt(WY) – THI with wild type levels of tyrosine and tryptophan

