## Supplementary text_S1 for "Phenylketonuria: modelling cerebral amino acid and neurotransmitter metabolism"

### Text S4.1. Model description

### 1 Glossary

**Table S1. Glossary of all abbreviations used in the text**

| Abbreviation | Full name |
|---|---|
| 3-Mt, Mt | 3-methoxytyramine |
| 5-Hiaa, Hiaa | 5-Hydroxyindoleacetic acid |
| 5-Htp, Htp | 5-Hydroxytryptophan |
| AA | Amino acid |
| AADC | Aromatic L-amino acid decarboxylase |
| COMT | Catechol-O-methyltransferase |
| Da | Dopamine |
| DBH | Dopamine beta-hydroxylase |
| F, Phe | Phenylalanine |
| H, His | Histidine |
| Ht | Serotonin |
| I, Ile | Isoleucine |
| $K_{eq}$ | Equilibrium constant |
| $K_i$ | Inhibitory (dissociation) constant |
| $K_{ic}$ | Inhibitory (dissociation) constant, competitive inhibitor |
| $K_{inc}$ | Inhibitory (dissociation) constant, non-competitive inhibitor |
| $K_{is}$ | Inhibitory (dissociation) constant of a substrate |
| $K_m$ | The Michaelis-Menten constant |
| L, Leu | Leucine |
| L-dopa | L-3,4-dihydroxyphenylalanine |
| LAT1 | Large neutral amino acid transporter 1 |
| LNAA-$Na^+$ | $Na^+$-dependent large neutral amino acid transporter |
| MAO | Monoamine oxidase |
| MV | Total pool of methionine and valine |
| Ne | Norepinephrine |
| Nmn | Normetanephrine |
| Q, Gln | Glutamine |
| sc | Stoichiometric coefficient |
| sf | Specificity factor |
| T, Thr | Threonine |
| TPH2 | Tryptophan monooxygenase 2 |
| TH | Tyrosine 3-monooxygenase |
| $v$ | Net rate of the reaction |
| $V$ | Maximum velocity of an enzyme |
| W, Trp | Tryptophan |
| Y, Tyr | Tyrosine |
| $y^+$ | Cationic amino acid transporter, $y^+$ system |

### 2 Kinetic rate equations

In our model, the majority of enzymes catalyze the conversion of multiple substrates (see Fig.1). However, each enzyme is characterized by its unique rate equation, with variable, substrate-specific(indicated by the subscript), rats $v$. Furthermore, LAT1 and y+ transporters are present on both sides of the blood-brain barrier at different concentrations. To address this, we use L for the luminal side ('blood') and A for the abluminal side ('brain') in the rate equations. In the abbreviations CELL indicates metabolites in the endothelial cell compartment (blood-brain barrier), BR indicates brain concentrations, and BL indicates blood concentrations. Additionally, the abbreviation AA is used for a general amino acid, and MV for a combined pool of methionine (Met, M) and valine (Val, V) to simplify the equation nomenclature. Most of the equations used in the model are of (ir)reversible Michaelis-Menten type. Exceptions to this rule are the rate equations for the transporter LAT1,

and two hydroxylases: TH and TPH2.
The model for LAT1 transporter follows the ping-pong bi-bi kinetics. It is known, that for each amino acid, LAT 1 transporter displays different maximum velocities (specificity factor $sf_{AA}$). Therefore a mean of sf factors was used for each exchange reaction. Additionally, LAT1 is known to have 2 x higher abundance on the luminal side $sf_{latL}$ of the blood-brain-barrier than on the abluminal side $sf_{latA}$, which was reflected in the rate equation.
Tyrosine and tryptophan hydroxylases are known to follow non-reversible Michaelis-Menten kinetics with substrate [1] and product inhibition.
The unit for the rates in the model description is $\mu mol \cdot min^{-1} \cdot mouse^{-1}$. The rate equations for the sink reactions, and consumption of the end products 3-Mt, Nmn, and 5-Hiaa, were constructed so that the sink reactions do not control the flux.

$$v_{lat1L(AA1)(AA2)} = \frac{V_{LAT1} \cdot sf_{\mathrm{latL}} \cdot \frac{sf_{\mathrm{AA1}}+sf_{\mathrm{AA2}}}{2} \cdot \left(\frac{\mathrm{AA1_{BL}[t] \cdot AA2_{CELL}[t]}}{Km_{\mathrm{AA1}} \cdot Km_{\mathrm{AA2}}} - \frac{\mathrm{AA1_{CELL}[t] \cdot AA2_{BL}[t]}}{Km_{\mathrm{AA1}} \cdot Km_{\mathrm{AA2}} \cdot K_{eq}}\right)}{\mathrm{subUnit1 \cdot subUnit2}}$$

where:

$$\begin{aligned} & AA \rightarrow Phe, Tyr, Trp, His, Ile, Leu, Thr, Gln, MV \\ & \mathrm{subUnit1} = 1 + \frac{\mathrm{AA1_{BL}[t]}}{Km_{\mathrm{AA1}}} + \frac{\mathrm{AA2_{BL}[t]}}{Km_{\mathrm{AA2}}} + \frac{\mathrm{AA3_{BL}[t]}}{Km_{\mathrm{AA3}}} + \frac{\mathrm{AA4_{BL}[t]}}{Km_{\mathrm{AA4}}} + \frac{\mathrm{AA5_{BL}[t]}}{Km_{\mathrm{AA5}}} + \frac{\mathrm{AA6_{BL}[t]}}{Km_{\mathrm{AA6}}} + \frac{\mathrm{AA7_{BL}[t]}}{Km_{\mathrm{AA7}}} + \frac{\mathrm{AA8_{BL}[t]}}{Km_{\mathrm{AA8}}} \\ & \quad + \frac{\mathrm{AA9_{BL}[t]}}{Km_{\mathrm{AA9}}} \\ & \mathrm{subUnit2} = 1 + \frac{\mathrm{AA1_{CELL}[t]}}{Km_{\mathrm{AA1}}} + \frac{\mathrm{AA2_{CELL}[t]}}{Km_{\mathrm{AA2}}} + \frac{\mathrm{AA3_{CELL}[t]}}{Km_{\mathrm{AA3}}} + \frac{\mathrm{AA4_{CELL}[t]}}{Km_{\mathrm{AA4}}} + \frac{\mathrm{AA5_{CELL}[t]}}{Km_{\mathrm{AA5}}} \frac{\mathrm{AA6_{CELL}[t]}}{Km_{\mathrm{AA6[t]}}} + \frac{\mathrm{AA7_{CELL}[t]}}{Km_{\mathrm{AA7}}} \\ & \quad + \frac{\mathrm{AA8_{CELL}[t]}}{Km_{\mathrm{AA8}}} + \frac{\mathrm{AA9_{CELL}[t]}}{Km_{\mathrm{AA9}}} \end{aligned} \tag{1}$$

$$v_{lat1A(AA1)(AA2)} = \frac{V_{LAT1} \cdot sf_{\mathrm{latA}} \cdot \frac{sf_{\mathrm{AA1}}+sf_{\mathrm{AA2}}}{2} \cdot \left(\frac{\mathrm{AA1_{CELL}[t] \cdot AA2_{BR}[t]}}{Km_{\mathrm{AA1}} \cdot Km_{\mathrm{AA2}}} - \frac{\mathrm{AA1_{BR}[t] \cdot AA2_{CELL}[t]}}{Km_{\mathrm{AA1}} \cdot Km_{\mathrm{AA2}} \cdot K_{eq}}\right)}{\mathrm{subUnit1 \cdot subUnit2}}$$

where:

$$\begin{aligned} & AA \rightarrow Phe, Tyr, Trp, His, Ile, Leu, Thr, Gln, MV \\ & \mathrm{subUnit1} = 1 + \frac{\mathrm{AA1_{CELL}[t]}}{Km_{\mathrm{AA1}}} + \frac{\mathrm{AA2_{CELL}[t]}}{Km_{\mathrm{AA2}}} + \frac{\mathrm{AA3_{CELL}[t]}}{Km_{\mathrm{AA3}}} + \frac{\mathrm{AA4_{CELL}[t]}}{Km_{\mathrm{AA4}}} + \frac{\mathrm{AA5_{CELL}[t]}}{Km_{\mathrm{AA5}}} + \frac{\mathrm{AA6_{CELL}[t]}}{Km_{\mathrm{AA6}}} \\ & \quad + \frac{\mathrm{AA7_{CELL}[t]}}{Km_{\mathrm{AA7}}} + \frac{\mathrm{AA8_{CELL}[t]}}{Km_{\mathrm{AA8}}} + \frac{\mathrm{AA9_{CELL}[t]}}{Km_{\mathrm{AA9}}} \\ & \mathrm{subUnit2} = 1 + \frac{\mathrm{AA1_{BR}[t]}}{Km_{\mathrm{AA1}}} + \frac{\mathrm{AA2_{BR}[t]}}{Km_{\mathrm{AA2}}} + \frac{\mathrm{AA3_{BR}[t]}}{Km_{\mathrm{AA3}}} + \frac{\mathrm{AA4_{BR}[t]}}{Km_{\mathrm{AA4}}} + \frac{\mathrm{AA5_{BR}[t]}}{Km_{\mathrm{AA5}}} \frac{\mathrm{AA6_{BR}[t]}}{Km_{\mathrm{AA6}}} + \frac{\mathrm{AA7_{BR}[t]}}{Km_{\mathrm{AA7}}} + \frac{\mathrm{AA8_{BR}[t]}}{Km_{\mathrm{AA8}}} \\ & \quad + \frac{\mathrm{AA9_{BR}[t]}}{Km_{\mathrm{AA9}}} \end{aligned} \tag{2}$$

$$v_{yL(AA1)} = \frac{Vy \cdot sf_{yL} \cdot \left(\frac{\mathrm{AA1_{BL}}}{Km_{\mathrm{AA1}}} - \frac{\mathrm{AA1_{CELL}}}{Km_{\mathrm{AA1}} \cdot Keq}\right)}{\left(1 + \frac{\mathrm{AA2_{BL}+AA2_{CELL}}}{Km_{\mathrm{AA1}}} + \frac{\mathrm{AA1_{BL}+AA1_{CELL}}}{Km_{\mathrm{AA2}}} + \frac{\mathrm{AA3_{BL}+AA3_{CELL}}}{Km_{\mathrm{AA3}}} + \frac{\mathrm{AA4_{BL}+AA4_{CELL}}}{Km_{\mathrm{AA4}}} + \frac{\mathrm{A5_{BL}+AA5_{CELL}}}{Km_{\mathrm{AA5}}}\right)}$$

where:

$$\begin{aligned} & AA \rightarrow Phe, His, Thr, Gln, MV \\ & Keq = 1 \end{aligned} \tag{3}$$

$$v_{yA(AA1)} = \frac{Vy \cdot sf_{yA} \cdot \left(\frac{\mathrm{AA1_{CELL}}}{Km_{\mathrm{AA1}}} - \frac{\mathrm{AA1_{BR}}}{Km_{\mathrm{AA1}} \cdot Keq}\right)}{\left(1 + \frac{\mathrm{AA2_{CELL}+AA2_{BR}}}{Km_{\mathrm{AA1}}} + \frac{\mathrm{AA1_{CELL}+AA1_{BR}}}{Km_{\mathrm{AA2}}} + \frac{\mathrm{AA3_{CELL}+AA3_{BR}}}{Km_{\mathrm{AA3}}} + \frac{\mathrm{AA4_{CELL}+AA4_{BR}}}{Km_{\mathrm{AA4}}} + \frac{\mathrm{A5_{CELL}+AA5_{BR}}}{Km_{\mathrm{AA5}}}\right)}$$

where:

$$\begin{aligned} & AA \rightarrow Phe, His, Thr, Gln, MV \\ & Keq = 1 \end{aligned} \tag{4}$$

$$v_{lnaa(AA1)} = \frac{V_{lnaa} \cdot \frac{\mathrm{AA1_{BR}[t]}}{Km_{\mathrm{AA1}}}}{1 + \frac{\mathrm{AA1_{BR}[t]}}{Km_{\mathrm{AA1}}} + \frac{\mathrm{AA2_{BR}[t]}}{Km_{\mathrm{AA2}}} + \frac{\mathrm{AA3_{BR}[t]}}{Km_{\mathrm{AA3}}} + \frac{\mathrm{AA4_{BR}[t]}}{Km_{\mathrm{AA4}}} + \frac{\mathrm{AA5_{BR}[t]}}{Km_{\mathrm{AA5}}} + \frac{\mathrm{AA6_{BR}[t]}}{Km_{\mathrm{AA6}}} + \frac{\mathrm{AA7_{BR}[t]}}{Km_{\mathrm{AA7}}} + \frac{\mathrm{AA8_{BR}[t]}}{Km_{\mathrm{AA8}}}} \tag{5}$$

where:

$$AA \rightarrow Phe, Tyr, Trp, His, Ile, Leu, Gln, MV$$

$$v_{TH(Tyr)} = \frac{V_{TH} \cdot \frac{\mathrm{Tyr_{BR}[t]}}{Km_{\mathrm{Tyr}}}}{Den}$$

where: (6)

$$\mathrm{Den} = 1+\frac{\mathrm{Phe_{BR}[t]}}{Km_{\mathrm{Phe}}}+\frac{\mathrm{Ne_{BR}[t]}}{Km_{\mathrm{Ne}}}+\frac{\mathrm{lDopa_{BR}[t]}}{Km_{\mathrm{lDopa}}}+\frac{\mathrm{Da_{BR}[t]}}{Km_{\mathrm{Da}}}+\mathrm{Tyr_{BR}}\cdot\frac{1+\frac{\mathrm{Phe_{BR}[t]}}{Kinc_{\mathrm{Phe}}}}{Km_{Tyr}}$$
$$+\mathrm{Tyr_{BR}[t]}^2\cdot\frac{1+\frac{\mathrm{Phe_{BR}[t]}}{Km_{\mathrm{Phe}}}+\frac{\mathrm{Ne_{BR}[t]}}{Km_{\mathrm{Ne}}}+\frac{\mathrm{lDopa_{BR}[t]}}{Km_{\mathrm{lDopa}}}+\frac{\mathrm{Da_{BR}[t]}}{Km_{\mathrm{Da}}}}{Km_{\mathrm{Tyr}}\cdot Kis_{\mathrm{Tyr}}}$$

$$v_{TH(Phe)} = \frac{V_{TH} \cdot \frac{\mathrm{Phe_{BR}[t]}}{Km_{\mathrm{Phe}}}}{Den}$$

where: (7)

$$\mathrm{Den} = 1+\frac{\mathrm{Tyr_{BR}[t]}}{Km_{\mathrm{Tyr}}}+\frac{\mathrm{Ne_{BR}[t]}}{Km_{\mathrm{Ne}}}+\frac{\mathrm{lDopa_{BR}[t]}}{Km_{\mathrm{lDopa}}}+\frac{\mathrm{Da_{BR}[t]}}{Km_{\mathrm{Da}}}+\mathrm{Phe_{BR}}\cdot\frac{1+\frac{\mathrm{Phe_{BR}[t]}}{Kinc_{\mathrm{Phe}}}}{Km_{Phe}}$$
$$+\mathrm{Phe_{BR}[t]}^2\cdot\frac{1+\frac{\mathrm{Tyr_{BR}[t]}}{Km_{\mathrm{Tyr}}}+\frac{\mathrm{Ne_{BR}[t]}}{Km_{\mathrm{Ne}}}+\frac{\mathrm{lDopa_{BR}[t]}}{Km_{\mathrm{lDopa}}}+\frac{\mathrm{Da_{BR}[t]}}{Km_{\mathrm{Da}}}}{Km_{\mathrm{Phe}}\cdot Kis_{\mathrm{Phe}}}$$

$$v_{TPH(Trp)} = \frac{V_{TPH} \cdot \frac{\mathrm{Trp_{BR}[t]}}{Km_{\mathrm{Trp}}}}{Den}$$

where: (8)

$$\mathrm{Den} = 1+\frac{\mathrm{Phe_{BR}[t]}}{Km_{\mathrm{Phe}}}+\frac{\mathrm{Htp_{BR}[t]}}{Km_{\mathrm{Htp}}}+\frac{\mathrm{lDopa_{BR}[t]}}{Km_{\mathrm{lDopa}}}+\frac{\mathrm{Da_{BR}[t]}}{Km_{\mathrm{Da}}}+\mathrm{Trp_{BR}}\cdot\frac{1+\frac{\mathrm{Phe_{BR}[t]}}{Kinc_{\mathrm{Phe}}}}{Km_{Trp}}$$
$$+\mathrm{Trp_{BR}[t]}^2\cdot\frac{1+\frac{\mathrm{Phe_{BR}[t]}}{Km_{\mathrm{Phe}}}+\frac{\mathrm{Htp_{BR}[t]}}{Km_{\mathrm{Htp}}}+\frac{\mathrm{lDopa_{BR}[t]}}{Km_{\mathrm{lDopa}}}+\frac{\mathrm{Da_{BR}[t]}}{Km_{\mathrm{Da}}}}{Km_{\mathrm{Trp}}\cdot Kis_{\mathrm{Trp}}}$$

$$v_{TPH(Phe)} = \frac{V_{TPH} \cdot \frac{\mathrm{Phe_{BR}[t]}}{Km_{\mathrm{Phe}}}}{Den}$$

where: (9)

$$\mathrm{Den} = 1+\frac{\mathrm{Trp_{BR}[t]}}{Km_{\mathrm{Trp}}}+\frac{\mathrm{Htp_{BR}[t]}}{Km_{\mathrm{Htp}}}+\frac{\mathrm{lDopa_{BR}[t]}}{Km_{\mathrm{lDopa}}}+\frac{\mathrm{Da_{BR}[t]}}{Km_{\mathrm{Da}}}+\mathrm{Phe_{BR}}\cdot\frac{1+\frac{\mathrm{Phe_{BR}[t]}}{Kinc_{\mathrm{Phe}}}}{Km_{Phe}}$$
$$+\mathrm{Phe_{BR}[t]}^2\cdot\frac{1+\frac{\mathrm{Trp_{BR}[t]}}{Km_{\mathrm{Trp}}}+\frac{\mathrm{Htp_{BR}[t]}}{Km_{\mathrm{Htp}}}+\frac{\mathrm{lDopa_{BR}[t]}}{Km_{\mathrm{lDopa}}}+\frac{\mathrm{Da_{BR}[t]}}{Km_{\mathrm{Da}}}}{Km_{\mathrm{Phe}}\cdot Kis_{\mathrm{Phe}}}$$

$$v_{AADC(s)\ (s\rightarrow lDopa|Htp)} = \frac{V_{AADC} \cdot sf_s \cdot \frac{\mathrm{S1[t]}}{Km_{\mathrm{S1}}}}{1+\frac{\mathrm{S1[t]}}{Km_{\mathrm{S1}}}+\frac{\mathrm{S2[t]}}{Km_{\mathrm{S2}}}} \tag{10}$$

$$v_{COMT(s)\ (s\rightarrow Da|Ne)} = \frac{V_{COMT} \cdot \frac{\mathrm{S1[t]}}{Km_{\mathrm{S1}}}}{1+\frac{\mathrm{S1[t]}}{Km_{\mathrm{S1}}}+\frac{\mathrm{S2[t]}}{Km_{\mathrm{S2}}}} \tag{11}$$

$$v_{DBH} = \frac{V_{DBH} \cdot \frac{\mathrm{Da[t]}}{Km_{\mathrm{Da}}}}{1+\frac{\mathrm{Da_{BR}[t]}}{Km_{\mathrm{Da}}}+\frac{\mathrm{Ne_{BR}[t]}}{Km_{\mathrm{Ne}}}+\frac{\mathrm{His_{BR}[t]}}{Km_{\mathrm{His}}}} \tag{12}$$

$$v_{MAO} = \frac{V_{MAO} \cdot \frac{\mathrm{Ht[t]}}{Km_{\mathrm{Ht}}}}{1+\frac{\mathrm{Ht[t]}}{Km_{\mathrm{Ht}}}} \tag{13}$$

$$sink_{S\ (S \to 3-Mt, Nmn, 5-Hiaa)} = Ks_S \cdot \frac{S[t]}{Km_S} \tag{14}$$

$$\begin{aligned} protSynB = Ks_{prot} \cdot & \frac{\mathrm{AA1_{BR}}}{\mathrm{AA1_{BR}} + \mathrm{Km_{AA1}}} \cdot \frac{\mathrm{AA2_{BR}}}{\mathrm{AA2_{BR}} + \mathrm{Km_{AA2}}} \cdot \frac{\mathrm{AA3_{BR}}}{\mathrm{AA3_{BR}} + \mathrm{Km_{AA3}}} \cdot \frac{\mathrm{AA4_{BR}}}{\mathrm{AA4_{BR}} + \mathrm{Km_{AA4}}} \cdot \\ & \frac{\mathrm{AA5_{BR}}}{\mathrm{AA5_{BR}} + \mathrm{Km_{AA5}}} \cdot \frac{\mathrm{AA6_{BR}}}{\mathrm{AA6_{BR}} + \mathrm{Km_{AA6}}} \cdot \frac{\mathrm{AA7_{BR}}}{\mathrm{AA7_{BR}} + \mathrm{Km_{AA7}}} \cdot \frac{\mathrm{AA8_{BR}}}{\mathrm{AA8_{BR}} + \mathrm{Km_{AA8}}} \cdot \\ & \frac{\mathrm{AA9_{BR}}}{\mathrm{AA9_{BR}} + \mathrm{Km_{AA9}}} \\ & \text{where:} \\ & {}_{AA \to Phe, Tyr, Trp, His, Ile, Leu, Thr, Gln, MV} \end{aligned} \tag{15}$$

$$\begin{aligned} protSynC = Ks_{prot} \cdot & \frac{\mathrm{AA1_{CELL}}}{\mathrm{AA1_{CELL}} + \mathrm{Km_{AA1}}} \cdot \frac{\mathrm{AA2_{CELL}}}{\mathrm{AA2_{BR}} + \mathrm{Km_{AA2}}} \cdot \frac{\mathrm{AA3_{CELL}}}{\mathrm{AA3_{v}} + \mathrm{Km_{AA3}}} \cdot \frac{\mathrm{AA4_{CELL}}}{\mathrm{AA4_{CELL}} + \mathrm{Km_{AA4}}} \cdot \\ & \frac{\mathrm{AA5_{CELL}}}{\mathrm{AA5_{CELL}} + \mathrm{Km_{AA5}}} \cdot \frac{\mathrm{AA6_{CELL}}}{\mathrm{AA6_{v}} + \mathrm{Km_{AA6}}} \cdot \frac{\mathrm{AA7_{CELL}}}{\mathrm{AA7_{CELL}} + \mathrm{Km_{AA7}}} \cdot \frac{\mathrm{AA8_{CELL}}}{\mathrm{AA8_{CELL}} + \mathrm{Km_{AA8}}} \cdot \\ & \frac{\mathrm{AA9_{CELL}}}{\mathrm{AA9_{CELL}} + \mathrm{Km_{AA9}}} \\ & \text{where:} \\ & {}_{AA \to Phe, Tyr, Trp, His, Ile, Leu, Thr, Gln, MV} \end{aligned} \tag{16}$$

### 3 Ordinary differential equations

Based on the reaction scheme in Figure 1, a set of 26 ordinary differential equations was created. As seen in Fig.1, many enzymes/transporters have broad substrate specificity, and many substrates can be converted by different enzymes/transporters. Furthermore, some transporters are localized in two different membranes, luminal (L) and abluminal(A). For instance, $v_{lat1LPheTyr}$ is the net rate of the luminal (L) transport of the Phe (phenylalanine) from the blood to the cell compartment, in exchange for the Tyr (tyrosine) which is transported by the LAT1 transporter in the opposite direction. In the abbreviations CELL indicates endothelial cell metabolite pool (blood-brain barrier), BR indicates total brain metabolite pool.

$$\begin{aligned} \frac{d\mathrm{PheCELL} \cdot V_{\mathrm{CELL}}}{dt} = & \, v_{lat1LPheTyr} + v_{lat1LPheTrp} + v_{lat1LPheHis} + v_{lat1LPheGln} + v_{lat1LPheIle} \\ & + v_{lat1LPheLeu} + v_{lat1LPheThr} + v_{lat1LPheMV} + v_{lnaaPhe} + v_{yLPhe} \\ & - v_{lat1APheTyr} - v_{lat1APheTrp} - v_{lat1APheHis} - v_{lat1APheGln} - v_{lat1APheIle} \\ & - v_{lat1APheLeu} - v_{lat1APheThr} - v_{lat1APheMV} - v_{yAPhe} - sc_{Phe} \cdot v_{ProtSynC} \end{aligned} \tag{17}$$

$$\begin{aligned} \frac{d\mathrm{TyrCELL} \cdot V_{\mathrm{CELL}}}{dt} = & \, v_{lat1APheTyr} + v_{lat1LTyrTrp} + v_{lat1LTyrHis} + v_{lat1LTyrGln} + v_{lat1LTyrIle} \\ & + v_{lat1LTyrLeu} + v_{lat1LTyrThr} + v_{lat1LTyrMV} + v_{lnaaTyr} - v_{lat1LPheTyr} \\ & - v_{lat1ATyrTrp} - v_{lat1ATyrHis} - v_{lat1ATyrGln} - v_{lat1ATyrIle} - v_{lat1ATyrLeu} \\ & - v_{lat1ATyrThr} - v_{lat1ATyrMV} - sc_{Tyr} \cdot v_{ProtSynC} \end{aligned} \tag{18}$$

$$\begin{aligned} \frac{d\mathrm{TrpCELL} \cdot V_{\mathrm{CELL}}}{dt} = & \, v_{lat1LTrpHis} + v_{lat1LTrpGln} + v_{lat1LTrpIle} + v_{lat1LTrpLeu} + v_{lat1LTrpThr} \\ & + v_{lat1LTrpMV} + v_{lat1APheTrp} + v_{lat1ATyrTrp} + v_{lnaaTrp} - v_{lat1LPheTrp} \\ & - v_{lat1LTyrTrp} - v_{lat1ATrpHis} - v_{lat1ATrpGln} - v_{lat1ATrpIle} - v_{lat1ATrpLeu} \\ & - v_{lat1ATrpThr} - v_{lat1ATrpMV} - sc_{Trp} \cdot v_{ProtSynC} \end{aligned} \tag{19}$$

$$\frac{d\text{HisCELL} \cdot V_{\text{CELL}}}{dt} = v_{lat1LHisGln} + v_{lat1LHisIle} + v_{lat1LHisLeu} + v_{lat1LHisThr} + v_{lat1LHisMV} + v_{lat1APheHis} + v_{lat1ATyrHis} + v_{lat1ATrpHis} + v_{lnaaHis} + v_{yLHis} - v_{lat1LPheHis} - v_{lat1LTyrHis} - v_{lat1LTrpHis} - v_{lat1AHisGln} - v_{lat1AHisIle} - v_{lat1AHisLeu} - v_{lat1AHisThr} - v_{lat1AHisMV} - v_{yAHis} - sc_{His} \cdot v_{ProtSynC} \tag{20}$$

$$\frac{d\text{IleCELL} \cdot V_{\text{CELL}}}{dt} = v_{lat1LIleLeu} + v_{lat1LIleThr} + v_{lat1LIleMV} + v_{lat1APheIle} + v_{lat1ATyrIle} + v_{lat1ATrpIle} + v_{lat1AHisIle} + v_{lat1AGlnIle} + v_{lnaaIle} - v_{lat1LPheIle} - v_{lat1LTyrIle} - v_{lat1LTrpIle} - v_{lat1LHisIle} - v_{lat1LGlnIle} - v_{lat1AIleLeu} - v_{lat1AIleThr} - v_{lat1AIleMV} - sc_{Ile} \cdot v_{ProtSynC} \tag{21}$$

$$\frac{d\text{LeuCELL} \cdot V_{\text{CELL}}}{dt} = v_{lat1LLeuThr} + v_{lat1LLeuMV} + v_{lat1APheLeu} + v_{lat1ATyrLeu} + v_{lat1ATrpLeu} + v_{lat1AHisLeu} + v_{lat1AGlnLeu} + v_{lat1AIleLeu} + v_{lnaaLeu} - v_{lat1LPheLeu} - v_{lat1LTyrLeu} - v_{lat1LTrpLeu} - v_{lat1LHisLeu} - v_{lat1LGlnLeu} - v_{lat1LIleLeu} - v_{lat1ALeuThr} - v_{lat1ALeuMV} - sc_{Leu} \cdot v_{ProtSynC} \tag{22}$$

$$\frac{d\text{ThrCELL} \cdot V_{\text{CELL}}}{dt} = v_{lat1LThrMV} + v_{lat1APheThr} + v_{lat1ATyrThr} + v_{lat1ATrpThr} + v_{lat1AHisThr} + v_{lat1AGlnThr} + v_{lat1AIleThr} + v_{lat1ALeuThr} + v_{lnaaThr} + v_{yLThr} - v_{lat1LPheThr} - v_{lat1LTyrThr} - v_{lat1LTrpThr} - v_{lat1LHisThr} - v_{lat1LGlnThr} - v_{lat1LIleThr} - v_{lat1LLeuThr} - v_{lat1AThrMV} - v_{yAThr} - sc_{Thr} \cdot v_{ProtSynC} \tag{23}$$

$$\frac{d\text{GlnCELL} \cdot V_{\text{CELL}}}{dt} = v_{lat1LGlnIle} + v_{lat1LGlnLeu} + v_{lat1LGlnThr} + v_{lat1LGlnMV} + v_{lat1APheGln} + v_{lat1ATyrGln} + v_{lat1ATrpGln} + v_{lat1AHisGln} + v_{yLGln} - v_{lat1LPheGln} - v_{lat1LTyrGln} - v_{lat1LTrpGln} - v_{lat1LHisGln} - v_{lat1AGlnIle} - v_{lat1AGlnLeu} - v_{lat1AGlnThr} - v_{lat1AGlnMV} - v_{yAGln} - sc_{Gln} \cdot v_{ProtSynC} \tag{24}$$

$$\frac{d\text{MVCELL} \cdot V_{\text{CELL}}}{dt} = v_{lat1APheMV} + v_{lat1ATyrMV} + v_{lat1ATrpMV} + v_{lat1AHisMV} + v_{lat1AGlnMV} + v_{lat1AIleMV} + v_{lat1ALeuMV} + v_{lat1AThrMV} + v_{lnaaMV} + v_{yLMV} - v_{lat1LPheMV} - v_{lat1LTyrMV} - v_{lat1LTrpMV} - v_{lat1LHisMV} - v_{lat1LGlnMV} - v_{lat1LIleMV} - v_{lat1LLeuMV} - v_{lat1LThrMV} - v_{yAMV} - sc_{MV} \cdot v_{ProtSynC} \tag{25}$$

$$\frac{d\text{PheBR} \cdot V_{\text{BR}}}{dt} = v_{lat1APheTyr} + v_{lat1APheTrp} + v_{lat1APheHis} + v_{lat1APheGln} + v_{lat1APheIle} + v_{lat1APheLeu} + v_{lat1APheThr} + v_{lat1APheMV} + v_{yAPhe} - v_{lnaaPhe} - v_{THPhe} - v_{TPHPhe} - sc_{Phe} \cdot v_{ProtSynB} \tag{26}$$

$$\frac{d\text{TyrBR} \cdot V_{\text{BR}}}{dt} = v_{lat1ATyrTrp} + v_{lat1ATyrHis} + v_{lat1ATyrGln} + v_{lat1ATyrIle} + v_{lat1ATyrLeu} + v_{lat1ATyrThr} + v_{lat1ATyrMV} + v_{TPHPhe} - v_{lat1APheTyr} - v_{lnaaTyr} - v_{THTyr} - sc_{Tyr} \cdot v_{ProtSyn} \tag{27}$$

$$\frac{d\text{TrpBR} \cdot V_{\text{BR}}}{dt} = v_{lat1ATrpHis} + v_{lat1ATrpGln} + v_{lat1ATrpIle} + v_{lat1ATrpLeu} + v_{lat1ATrpThr} + v_{lat1ATrpMV} - v_{lat1APheTrp} - v_{lat1ATyrTrp} - v_{lnaaTrp} - v_{TPHTrp} - sc_{Trp} \cdot v_{ProtSyn} \tag{28}$$

$$\frac{d\text{HisBR}\cdot V_{\text{BR}}}{dt} = v_{lat1AHisGln} + v_{lat1AHisIle} + v_{lat1AHisLeu} + v_{lat1AHisThr} + v_{lat1AHisMV} + v_{yAHis} - v_{lat1APheHis} - v_{lat1ATyrHis} - v_{lat1ATrpHis} - v_{lnaaHis} - sc_{His}\cdot v_{ProtSyn} \tag{29}$$

$$\frac{d\text{IleBR}\cdot V_{\text{BR}}}{dt} = v_{lat1AIleLeu} + v_{lat1AIleThr} + v_{lat1AIleMV} - v_{lat1APheIle} - v_{lat1ATyrIle} - v_{lat1ATrpIle} - v_{lat1AHisIle} - v_{lat1AGlnIle} - v_{lnaaIle} - sc_{Ile}\cdot v_{ProtSyn} \tag{30}$$

$$\frac{d\text{LeuBR}\cdot V_{\text{BR}}}{dt} = v_{lat1ALeuThr} + v_{lat1ALeuMV} - v_{lat1APheLeu} - v_{lat1ATyrLeu} - v_{lat1ATrpLeu} - v_{lat1AHisLeu} - v_{lat1AGlnLeu} - v_{lat1AIleLeu} - v_{lnaaLeu} - sc_{Leu}\cdot v_{ProtSyn} \tag{31}$$

$$\frac{d\text{ThrBR}\cdot V_{\text{BRAIN}}}{dt} = v_{lat1AThrMV} + v_{yAThr} - v_{lat1APheThr} - v_{lat1ATyrThr} - v_{lat1ATrpThr} - v_{lat1AHisThr} - v_{lat1AGlnThr} - v_{lat1AIleThr} - v_{lat1ALeuThr} - v_{lnaaThr} - sc_{Thr}\cdot v_{ProtSyn} \tag{32}$$

$$\frac{d\text{GlnBR}\cdot V_{\text{BR}}}{dt} = v_{lat1AGlnIle} + v_{lat1AGlnLeu} + v_{lat1AGlnThr} + v_{lat1AGlnMV} + v_{yAGln} - v_{lat1APheGln} - v_{lat1ATyrGln} - v_{lat1ATrpGln} - v_{lat1AHisGln} - sc_{Gln}\cdot v_{ProtSyn} \tag{33}$$

$$\frac{d\text{MVBR}\cdot V_{\text{BR}}}{dt} = v_{yAMV} - v_{lat1APheMV} - v_{lat1ATyrMV} - v_{lat1ATrpMV} - v_{lat1AHisMV} - v_{lat1AGlnMV} - v_{lat1AIleMV} - v_{lat1ALeuMV} - v_{lat1AThrMV} - v_{lnaaMV} - sc_{MV}\cdot v_{ProtSyn} \tag{34}$$

$$\frac{d\text{Htp}\cdot V_{\text{BR}}}{dt} = v_{TPHTrp} - v_{AADChtp} \tag{35}$$

$$\frac{d\text{Ht}\cdot V_{\text{BR}}}{dt} = v_{AADChtp} - v_{MAO} \tag{36}$$

$$\frac{d\text{Hiaa}\cdot V_{\text{BR}}}{dt} = v_{MAO} - sinkHiaa \tag{37}$$

$$\frac{d\text{lDopa}\cdot V_{\text{BR}}}{dt} = v_{THTyr} + v_{THPhe} - v_{AADCdopa} \tag{38}$$

$$\frac{d\text{Da}\cdot V_{\text{BR}}}{dt} = v_{AADCdopa} - v_{DBH} - v_{COMTda} \tag{39}$$

$$\frac{d\text{Mt}\cdot V_{\text{BR}}}{dt} = v_{COMTda} - sink3Mt \tag{40}$$

$$\frac{d\text{Ne}\cdot V_{\text{BR}}}{dt} = v_{DBH} - v_{COMTne} \tag{41}$$

$$\frac{d\text{Nmn}\cdot V_{\text{BR}}}{dt} = v_{COMTne} - sinkNmn \tag{42}$$

### 4 Simulations of diets - model validation

Parameters used in the model are summarized in Table S2. For the y+ and LAT1 transporters, the equilibrium constants were assumed to be 1, as these are not driven by any external Gibbs-energy input. All maximum enzyme velocities from the literature were normalized to the total brain protein content (47.5 mg prot · brain$^{-1}$), taking into account different expression levels of each enzyme in the brain where needed. To this end, we calculated weighted means of transcript levels for each enzyme based on the differential expression levels in the mouse brain and then normalized them to LAT1 (see Table S3).

**Table S2. Kinetic parameters.**

| Parameter | Value | | Reference |
|---|---|---|---|
| **LAT1 transporter** | | | |
| $V_{LAT1}$ | 1.9665 | μmol · min$^{-1}$· mouse$^{-1}$ | fitted manually to the WT data |
| $sf_{Phe}$ | 1 | | [2] |
| $sf_{Tyr}$ | 2.341 | | [2] |
| $sf_{Trp}$ | 0.854 | | [2] |
| $sf_{His}$ | 1.488 | | [2] |
| $sf_{Gln}$ | 1.049 | | [2] |
| $sf_{Ile}$ | 1.463 | | [2] |
| $sf_{Leu}$ | 1.439 | | [2] |
| $sf_{Thr}$ | 0.415 | | [2] |
| $sf_{MV}$ | 1.042 | | [2] |
| $sf_{LatL}$ | 2 | | 2x higher expression on the luminal site [3] |
| $sf_{LatA}$ | 1 | | |
| $Keq$ | 1 | | |
| $Km_{Phe}$ | 11 | μM | [2] |
| $Km_{Tyr}$ | 64 | μM | [2] |
| $Km_{Trp}$ | 15 | μM | [2] |
| $Km_{His}$ | 100 | μM | [2] |
| $Km_{Gln}$ | 880 | μM | [2] |
| $Km_{Ile}$ | 56 | μM | [2] |
| $Km_{Leu}$ | 29 | μM | [2] |
| $Km_{Thr}$ | 220 | μM | [2] |
| $Km_{MV}$ | 165.59 | μM | weighted average for methionine and valine [2] (based on their abundance in the brain in WT mice) |
| **y+ transporter** | | | |
| $V_y$ | 0.1045 | μmol · min$^{-1}$· mouse$^{-1}$ | original value: 2.2 nmol · min$^{-1}$· mg prot.$^{-1}$, bovine [4] |
| $sf_{yL}$ | 1 | | 2x higher expression on the abluminal site [3] |
| $sf_{yA}$ | 2 | | |
| $Keq$ | 1 | | |
| $Km_{Phe}$ | 590 | μM | [4] |
| $Km_{His}$ | 630 | μM | [4] |
| $Km_{Gln}$ | 620 | μM | [4] |
| $Km_{Thr}$ | 670 | μM | [4] |
| $Km_{MV}$ | 687 | μM | [4] |
| **LNAA-Na$^+$ transporter** | | | |
| $V_{LNAA}$ | 0.005415 | μmol · min$^{-1}$· mouse$^{-1}$ | original value: 114 pmol · min$^{-1}$· mg prot$^{-1}$, bovine [5] |
| $Km_{Leu}$ | 21 | μM | [5] |
| $Km_{Phe}$ | 26.11 | μM | |
| $Km_{Tyr}$ | 25.87 | μM | |
| $Km_{Trp}$ | 24.33 | μM | rough estimates based on the reported changes in the apparent Vmax due to the substrate competition [5] |
| $Km_{His}$ | 30.57 | μM | |
| $Km_{Ile}$ | 21.32 | μM | |

**Table S2. Kinetic parameters (continued).**

| Parameter | Value | | Reference |
|---|---|---|---|
| $Km_{Thr}$ | 30.91 | $\mu$M | |
| $Km_{MV}$ | 27.00 | $\mu$M | |
| **TH** | | | |
| $V_{TH}$ | 0.001425 | $\mu$mol $\cdot$ min$^{-1}\cdot$ mouse$^{-1}$ | original value: 30 pmol $\cdot$ min$^{-1}\cdot$ mg prot$^{-1}$, rat midbrain slices [6] |
| $Km_{Tyr}$ | 17 | $\mu$M | [1] |
| $Kis_{Tyr}$ | 227 | $\mu$M | [1] |
| $Km_{Phe}$ | 103 | $\mu$M | [1] |
| $Kinc_{Phe}$ | 736 | $\mu$M | [1] |
| $Kis_{Phe}$ | 1375.3529 | $\mu$M | unknown, value based on the $^{Km_{Tyr}}/_{Kis_{Tyr}}$ ratio from [1] |
| $Kis_{Ne}$ | 280 | $\mu$M | [7] |
| $Kis_{Da}$ | 600 | $\mu$M | [7] |
| $Kis_{lDopa}$ | 56 | $\mu$M | [8] |
| **TPH2** | | | |
| $V_{TPH}$ | 0.00115 | $\mu$mol $\cdot$ min$^{-1}\cdot$ mouse$^{-1}$ | original value: 0.0242 nmol $\cdot$ min$^{-1}\cdot$ mg prot$^{-1}$, rat brain extract [9] |
| $Km_{Trp}$ | 13.2 | $\mu$M | [1] |
| $Kis_{Trp}$ | 1030 | $\mu$M | [1] |
| $Km_{Phe}$ | 72.7 | $\mu$M | [1] |
| $Kinc_{Phe}$ | 257 | $\mu$M | [1] |
| $Kis_{Phe}$ | 5672.8030 | $\mu$M | unknown, value based on the $^{Km_{Trp}}/_{Kis_{Trp}}$ ratio from [1] |
| $Kis_{Htp}$ | 35 | $\mu$M | [10] |
| $Kis_{Da}$ | 94 | $\mu$M | [11] |
| $Kis_{lDopa}$ | 17 | $\mu$M | [11] |
| **AADC** | | | |
| $V_{AADC}$ | 0.2831 | $\mu$mol $\cdot$ min$^{-1}\cdot$ mouse$^{-1}$ | original value: 5.96 nmol $\cdot$ min$^{-1}\cdot$ mg prot$^{-1}$, human [12] |
| $Sf_{lDopa}$ | 1 | $\mu$M | Vmax for l-Dopa is 23 times higher than |
| $Sf_{Htp}$ | 0.0435 | $\mu$M | for serotonin [13] |
| $Km_{lDopa}$ | 70 | $\mu$M | human, [14] |
| $Km_{Htp}$ | 47 | $\mu$M | human, [15] |
| **DBH** | | | |
| $V_{DBH}$ | 0.8102 | $\mu$mol $\cdot$ min$^{-1}\cdot$ mouse$^{-1}$ | original value: 1500 nmol $\cdot$ min$^{-1}\cdot$ mg prot$^{-1}$, Rat, purified protein [16], normalized to LAT1 |
| $Keq$ | $1\cdot10^{6}$ | $\mu$M | arbitrary number, the reaction is virtually irreversible, however, the product does slightly inhibit the enzyme |
| $Km_{Da}$ | 200 | $\mu$M | *Gallus gallus* [17] |
| $Km_{Ne}$ | 5000 | $\mu$M | *Gallus gallus* [17] |
| $Ki_{His}$ | 410 | $\mu$M | reported in BRENDA database after [18] |
| **MAO** | | | |
| $V_{MAO}$ | 0.1501 | $\mu$mol $\cdot$ min$^{-1}\cdot$ mouse$^{-1}$ | original value: 3.16 nmol $\cdot$ min$^{-1}\cdot$ mg prot$^{-1}$, Rat, cell lysates, at 100 $\mu$M serotonin [19] |
| $Km_{Ht}$ | 99 | $\mu$M | Rat [20] |
| **COMT** | | | |
| $V_{COMT}$ | 0.0494 | $\mu$mol $\cdot$ min$^{-1}\cdot$ mouse$^{-1}$ | original value: 1.04 nmol $\cdot$ min$^{-1}\cdot$ mg prot$^{-1}$, rat liver extract [21] |
| $Km_{Da}$ | 3.3 | $\mu$M | human [22] |
| $Km_{Ne}$ | 5.28 | $\mu$M | human, based on the ratio of $^{Km_{Ne}}/_{Km_{Da}}$ in recombinant Sf9 cells, and $Km_{Da}$ in human brain [22] |

**Table S2. Kinetic parameters (continued).**

| Parameter | Value | | Reference |
|---|---|---|---|
| **protSyn** | | | |
| $V_{protSyn}$ | 0.265 | $\mu$mol · min$^{-1}$· mouse$^{-1}$ | average rate based on the regional rates of cerebral Leu incorporation (5.4565 nmol · min$^{-1}$· g tissue$^{-1}$) [23] |
| $Km_{Phe}$ | 2.6 | $\mu$M | human [24] |
| $Km_{Tyr}$ | 34 | $\mu$M | human [25] |
| $Km_{Trp}$ | 7.4 | $\mu$M | human [26] |
| $Km_{His}$ | 30 | $\mu$M | *E.coli* [27] |
| $Km_{Gln}$ | 114 | $\mu$M | *E.coli* [28] |
| $Km_{Ile}$ | 52 | $\mu$M | *E.coli* [29] |
| $Km_{Leu}$ | 45.6 | $\mu$M | human [30] |
| $Km_{Thr}$ | 110 | $\mu$M | *E.coli* [31] |
| $Km_{MV}$ | 7.87877 | $\mu$M | Met: human [32], Val: *E.coli* [33] |
| $sc_{Phe}$ | 0.039 | | |
| $sc_{Tyr}$ | 0.029 | | |
| $sc_{Trp}$ | 0.012 | | |
| $sc_{His}$ | 0.026 | | protein composition based on the codon usage [34] |
| $sc_{Gln}$ | 0.046 | | |
| $sc_{Ile}$ | 0.045 | | |
| $sc_{Leu}$ | 0.105 | | |
| $sc_{Thr}$ | 0.055 | | |
| $sc_{MV}$ | 0.053 | | |
| **Sinks** | | | |
| $Ks_{Hiaa}$ | 30 | $\mu$mol · min$^{-1}$· mouse$^{-1}$ | |
| $Km_{Hiaa}$ | 49 | $\mu$M | |
| $Ks_{3-Mt}$ | 10 | $\mu$mol · min$^{-1}$· mouse$^{-1}$ | arbitrary values estimated to not display any metabolic control over the system |
| $Km_{3-Mt}$ | 50 | $\mu$M | |
| $Ks_{Nmn}$ | 20 | $\mu$mol · min$^{-1}$· mouse$^{-1}$ | |
| $Km_{Nmn}$ | 48 | $\mu$M | |
| **Volumes** | | | |
| $V_{BL}$ | 0.00126 | l · mouse$^{-1}$ | own data |
| $V_{CELL}$ | 7.47·10$^{-6}$ | l · mouse$^{-1}$ | surface area of microvessels: 150cm$^2$· g tissue$^-$1 [35]; brain mass = 475 mg [own data]; average endothelial cell volume: 1009$\mu$m$^3$ [36]; average area of ecapillary endothelial cell: 962 $\mu$m$^3$ [37] |
| $V_{BR}$ | 0.004532 | l · mouse$^{-1}$ | [38] |

**Table S3. Protein expression levels in the brain.**

| Protein | A | B | C | D | E | F | G | H | I | J | Weight. mean | Rel. to LAT1 |
|---|---|---|---|---|---|---|---|---|---|---|---|---|
| LAT1 | 19.50 | 22.58 | 23.45 | 33.86 | 17.60 | 18.05 | 21.34 | 27.94 | 13.80 | 29.33 | 24.89 | 1 |
| TPH2 | 0.01 | 0.03 | 0.05 | 0.02 | - | - | 0.03 | 0.03 | 0.01 | 0.01 | 0.03 | 0.001 |
| TH | - | - | - | - | - | - | - | 6.43 | - | 32.30 | 0.48 | 0.019 |
| AADC | - | - | - | - | - | - | - | 0.91 | - | 2.67 | 0.05 | 0.002 |
| DBH | - | - | - | 1.2 | - | - | - | - | - | - | 0.28 | 0.011 |
| MAO-A | 11.68 | 10.57 | 9.52 | 3.19 | 19.96 | 9.68 | 7.94 | 13.29 | 11.66 | 12.18 | 9.91 | 0.398 |
| COMT | 13.35 | 11.93 | 10.78 | 11.82 | 13.27 | 13.06 | 13.39 | 12.60 | 12.97 | 14.98 | 12.02 | 0.483 |
| LNAA | 1.92 | 3.13 | 4.33 | 2.94 | 2.20 | 1.99 | 2.17 | 4.67 | 2.03 | 2.41 | 3.16 | 0.127 |
| y+ | 4.85 | 8.76 | 9.81 | 9.64 | 4.45 | 3.73 | 6.69 | 8.64 | 4.03 | 6.28 | 8.34 | 0.335 |

A-Amygdala, B-Anterior cingulate cortex, C-Frontal cortex, D-Cortex, E-Caudate, F-Putamen, G-Hippocampus, H-Hypothalamus, I-Nucleus accumbens, J-Substantia nigra

The blood concentrations of amino acids were set, as fixed values, according to the measured values in mice (see Table S4). The initial concentrations (in $\mu M$) for the variables were first assigned arbitrarily, and then a time-course simulation was performed to acquire a set of initial concentrations close to the steady-state values for WT diet. This newly acquired set of initial concentrations was then used for all the simulations.

**Table S4. Average amino-acid concentrations in the blood, as used in the model (in $\mu M$ ).**

| Amino-acid | WT | PKU | LNAA (-Thr) | LNAA (+Thr) | Tyr+Trp | Leu+Ile | Thr | High protein |
|---|---|---|---|---|---|---|---|---|
| *Phe* | 304 | 1803 | 1387 | 1381 | 1886 | 1826 | 1848 | 2503 |
| *Tyr* | 131 | 61 | 119 | 96 | 172 | 61 | 59 | 86 |
| *Trp* | 147 | 98 | 172 | 177 | 168 | 104 | 105 | 89 |
| *His* | 79 | 68 | 107 | 87 | 63 | 66 | 65 | 72 |
| *Gln* | 604 | 487 | 464 | 507 | 511 | 420 | 494 | 399 |
| *Ile* | 148 | 120 | 273 | 213 | 114 | 358 | 112 | 180 |
| *Leu* | 221 | 167 | 303 | 246 | 151 | 470 | 153 | 243 |
| *Thr* | 314 | 208 | 202 | 484 | 182 | 198 | 311 | 227 |
| *MV* | 464 | 363 | 1052 | 752 | 325 | 410 | 330 | 539 |

WT - wild-type mice, standard chow; PKU - C57Bl/6 Pah-enu2 (PKU) mice, standard chow; LNAA(-Thr) - PKU mice, standard chow with LNAA(-Thr) supplementation; LNAA(+Thr) - PKU mice, standard chow with LNAA(+Thr) supplementation; Tyr+Trp - PKU mice, standard chow with Tyr and Trp supplementation; Leu+Ile - PKU mice, standard chow with Leu and Ile supplementation; Thr - PKU mice, standard chow with Thr supplementation; High protein - PKU mice, an isonitrogenic/isocaloric high-protein control diet

The simulation values represented in figures 2, 3, 4, 6, S2, S3, S9, and S12 were calculated as a weighted average of concentrations of amino acids in the BRAIN and CELL compartments as in the following equation:

$$AA_{total} = \frac{V_{CELL}}{V_{CELL} + V_{BR}} \cdot AA_{CELL} + \frac{V_{BR}}{V_{CELL} + V_{BR}} \cdot AA_{BR}$$

This approach reflects better the experimental data available, which does not distinguish between blood-brain-barrier and brain compartments.
